# A hypothalamic circuit for anticipating future changes in energy balance

**DOI:** 10.1101/2025.09.27.678865

**Authors:** Samuel J. Walker, Elijah D. Lowenstein, Amelia M. Douglass, Joseph C. Madara, Jon M. Resch, Jenkang Tao, Bradford B. Lowell

**Affiliations:** Division of Endocrinology, Diabetes and Metabolism, Department of Medicine, Beth Israel Deaconess Medical Center, Harvard Medical School, Boston, MA 02215, USA; Department of Neuroscience and Pharmacology, University of Iowa Carver College of Medicine, Iowa City, IA, USA; Program in Neuroscience, Harvard Medical School, Boston, MA 02115, USA

## Abstract

AgRP neurons cause hunger, the drive to seek and consume food. Their activation by fasting is key for survival and is thought to be triggered by feedback when energy stores are low. However, we know that environmental cues can also regulate AgRP neurons, since cues that predict future food intake rapidly inhibit AgRP neurons. But is the converse true: can the prediction of future fasting rapidly activate AgRP neurons? Here we show that such rapid fasting activation of AgRP neurons does occur. This fasting response is driven by excitatory input from paraventricular hypothalamic neurons expressing *Sim2*, which are bidirectionally sensitive to predictions of future energy state. In this way, cognitively-processed contextual information conveyed by PVH^Sim2^ neurons strongly activates AgRP neurons. Lastly, chronic silencing of PVH^Sim2^ neurons causes persistent hypophagia. This PVH^Sim2^ to AgRP neuron circuit, by anticipating and preventing negative energy balance, provides an important new dimension of hunger regulation.

## Introduction

The tight regulation of food intake is critical to fitness and survival. This regulation is achieved by a variety of neuron types in the brain, but how these neurons integrate long-timescale signals of systemic energy balance with shorter-timescale sensory and cognitive signals remains unclear.

Neurons in the arcuate nucleus of the hypothalamus that express the neuropeptide AgRP (ARC^AgRP^ neurons) are key players in energy balance regulation^1^. ARC^AgRP^ neurons have been termed “hunger neurons” since they show increased activity following fasting^2^, and experimentally increasing their activity recapitulates many behavioral, physiological, and neural consequences of fasting, including increased food intake and food-seeking behavior^1,3–6^. However, the signals that regulate ARC^AgRP^ neuron activity *in vivo* remain only partly understood.

During the process of eating-induced satiation, ARC^AgRP^ neurons integrate rapid feedforward sensory information with post-ingestive signals from the gut and slower, hormonal signals of fat stores, resulting in a rapid and sustained drop in their activity in response to food^7–12^. In contrast, how fasting increases the activity of ARC^AgRP^ neurons, and the timescale over which it does so, remains poorly understood. ARC^AgRP^ neurons are sensitive to circulating feedback signals of energy balance (e.g. leptin, insulin, ghrelin)^12–16^, and are thought to gradually increase their activity as energy stores are depleted^1,2^. However, challenging this view, recent long-term recordings suggest that ARC^AgRP^ neuron activity increases very early in fasting, before energy stores are depleted^17^. The origin of this rapid increase in ARC^AgRP^ neuron activity early in fasting is unknown.

Another key nucleus in the control of food intake is the paraventricular nucleus of the hypothalamus (PVH)^18^. The PVH is a multifunctional nucleus, containing neurons that regulate diverse autonomic, neuroendocrine, and behavioral processes, many related to energy balance^19,20^. In the context of food intake, the PVH has long been thought of as a *satiety center*: ablation of the PVH or inhibition of all PVH neurons increases food intake^21,22^, whereas activation of all PVH neurons inhibits feeding^23^. This influence on food intake is due to PVH “satiety neurons” that are downstream of inhibitory ARC^AgRP^ hunger neurons^2,18,24,25^.

Embedded within the PVH “satiety center”, however, is a subset of neurons with an opposing influence on food intake. Specifically, artificial activation of PVH neurons expressing *Trh* and *Adcyap1* increases food intake in sated mice^26^. These neurons increase food intake through their excitatory output onto ARC^AgRP^ neurons. Moreover, these PVH→ARC^AgRP^ connections undergo profound plasticity depending on energy state: negative energy balance increases the number of excitatory synaptic connections from PVH to ARC^AgRP^ neurons^27,28^. This plasticity is likely important for the regulation of food intake, since preventing it by deleting NMDA receptors from ARC^AgRP^ neurons reduces body weight and food intake^27^. However, the information that PVH^Trh/Adcyap1^ neurons transmit to ARC^AgRP^ neurons, and how they contribute to the physiological regulation of food intake, remains unclear^1^. This is in large part due to an inability to selectively access the PVH neurons that are afferent to ARC^AgRP^ neurons in order to record or chronically inhibit their activity, since both *Trh* and *Adcyap1* are expressed in a number of PVH cell types with diverse functions^19,29,30^.

In this study, we mined a transcriptomic atlas of PVH neurons to identify a specific PVH neuron type (PVH^Sim2^ neurons) that provides excitatory input to ARC^AgRP^ neurons and drives feeding. We demonstrate that these PVH^Sim2^ neurons transmit predictions of future energy balance to ARC^AgRP^ neurons. PVH^Sim2^ neurons are bidirectionally sensitive to food availability in the environment: food presentation inhibits PVH^Sim2^ neuron activity, whereas removal of food increases PVH^Sim2^ and ARC^AgRP^ neuron activity, likely as a response to unsuccessful food-seeking behavior. Finally, we show that this excitatory input to ARC^AgRP^ neurons is critical for long-term food intake regulation, since chronic inhibition of PVH^Sim2^ results in a chronic reduction in food intake.

## Results

### *Sim2* expression selectively labels *Trh/Pacap* neurons in the PVH

PVH neurons expressing *Trh* and *Adcyap1* provide monosynaptic input to ARC^AgRP^ neurons and regulate food intake^26,28^. However, each of these genes is expressed in multiple, functionally distinct PVH neuron types (for example, *Trh* is also expressed in neuroendocrine neurons that regulate thyroid hormone release^31^). This makes it difficult to gain selective genetic access to ARC^AgRP^ neuron afferents, thereby complicating efforts to selectively study their regulation and function. As a first step towards gaining selective experimental access to these neurons, we aimed to pinpoint the precise transcriptional identity of the PVH neurons that synapse onto ARC^AgRP^ neurons (Figure 1A). To do so, we examined the co-expression of *Trh* and *Adcyap1* in an unpublished sc/snRNAseq atlas of PVH neurons (Li, Butler *et al.*, unpublished). We specifically looked for neurons co-expressing both *Trh* and *Adcyap1*, because they are likely AgRP neuron afferents for the following reasons: *Trh+/Adcyap1+* co-expressing neurons are present in the PVH^26^, show increased expression of Fos following overnight fasting^28^, and increase food intake when activated (Figure S1).

**Figure 1.**
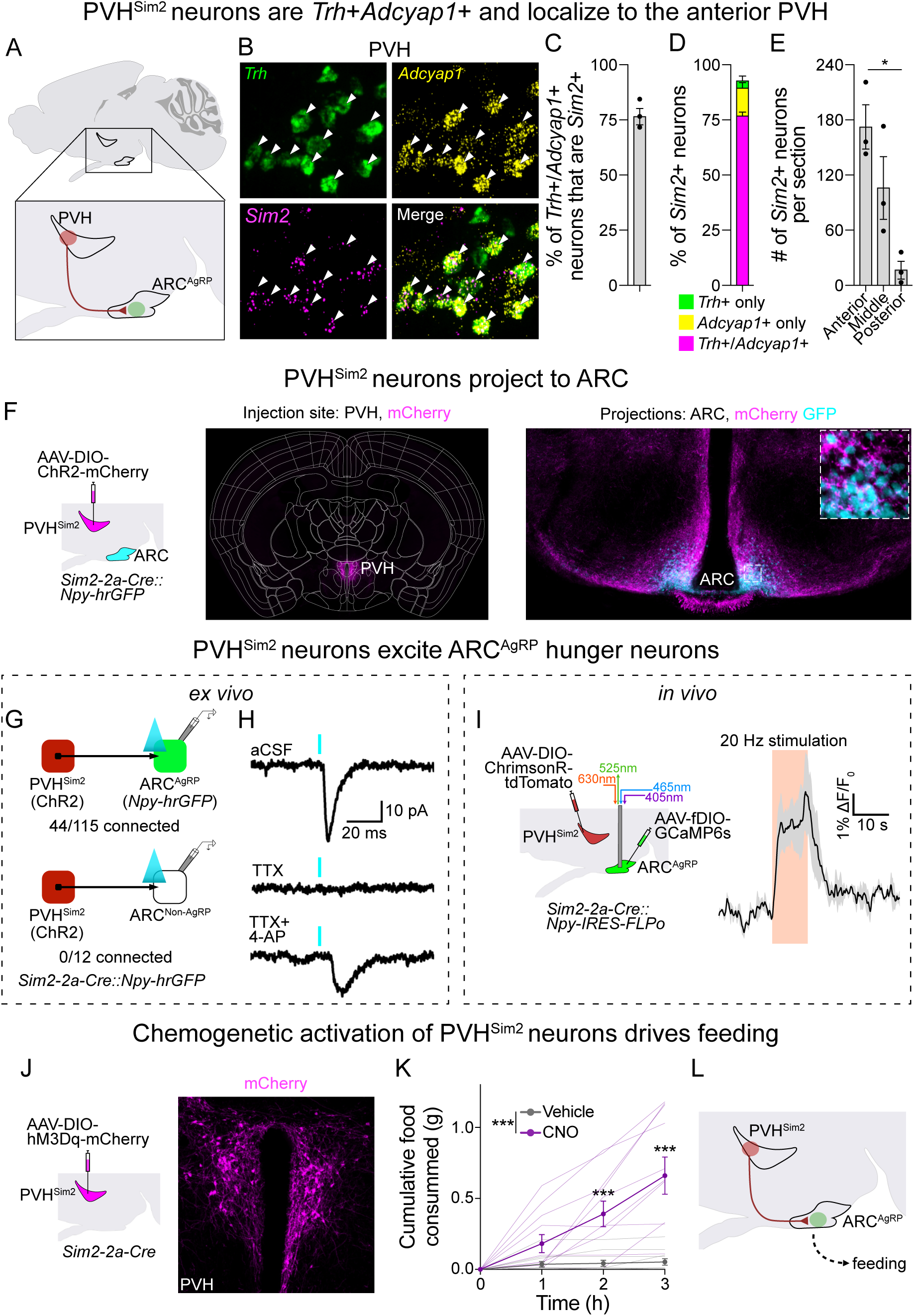
PVH^Sim2^ neurons excite ARC^AgRP^ neurons and drive feeding. A) Schematic of PVH neurons that synapse onto ARC^AgRP^ neurons. B) Representative image of *Trh* (green), *Adcyap1* (yellow) and *Sim2* (magenta) expression in the anterior PVH, as detected by single molecule fluorescent *in situ* hybridization (RNAscope). C) Percentage of *Trh*/*Adcyap1*-coexpressing PVH neurons that also express *Sim2*. (N = 3 mice, mean ± SEM). D) Percentage of *Sim2*-expressing PVH neurons that express *Trh* only, *Adcyap1* only, or both. (N = 3 mice, mean ± SEM). E) Number of *Sim2*-expressing neurons per section at different anterior-posterior levels of PVH. A: anterior; M: mid, P: posterior. (N = 3 mice, mean ± SEM; ordinary one-way ANOVA). F) Left: schematic of injection of AAV-DIO-ChR2-mCherry into the PVH of a *Sim2-2a-Cre*::*Npy-hrGFP* mouse. Middle: resultant mCherry expression in the PVH, and not surrounding areas. Right: mCherry-labelled PVH^Sim2^ neuron projections (magenta), and GFP-expressing ARC^AgRP^ neuron cell bodies (cyan), in the ARC. G) Fraction of Npy-hrGFP-positive (top) and –negative (bottom) ARC neurons that show leEPSCs with monosynaptic latency following optogenetic activation of PVH^Sim2^ neuron projections by CRACM. H) Example recording from Npy-hrGFP-positive neuron with optogenetic stimulation of PVH^Sim2^ neurons in the presence of artificial cerebrospinal fluid (aCSF) alone (top), or with tetrodotoxin (TTX, middle), or with TTX and 4-aminopyridine (4-AP, bottom). I) Left: schematic of *in vivo* optogenetic stimulation of PVH^Sim2^ terminals with simultaneous fiber photometry recording of ARC^AgRP^ neuron activity. Right: mean ± SEM trace of ARC^AgRP^ neuron activity before, during, and after 10s stimulation of PVH^Sim2^ axons in ARC. (N = 3 mice, mean over 15 stimulations per mouse). J) Left: schematic of AAV-DIO-hM3Dq-mCherry injection into the PVH of *Sim2-2a-Cre* mice. Right: representative image of resultant mCherry expression (magenta) in the PVH. K) Daytime (ZT3-6) food intake following injection of vehicle or CNO. Mean ± SEM; thin lines represent individual mice. (N = 11 mice; two-way repeated measures ANOVA; post-hoc comparisons at each time point with Sidak correction). L) Schematic of the PVH^Sim2^ to ARC^AgRP^ circuit and its role in driving feeding behavior. *p<0.05; **p<0.01; ***p<0.001.

We observed that the majority of *Trh+/Adcyap1+* neurons belonged to a transcriptional cluster that uniquely expressed the transcription factor *Sim2* (not to be confused with *Sim1*, which is broadly-expressed in the majority of PVH neurons^32^). We confirmed these results using three-color single molecule fluorescent *in situ* hybridization (smFISH) against *Trh*, *Adcyap1* and *Sim2* (Figure 1B). Indeed, throughout the PVH, 76.9 ± 3.7% (mean ± SEM) of *Trh+/Adcyap1+* neurons also expressed *Sim2* (Figure 1C). Similarly, 77.2 ± 1.8% of *Sim2*-expressing neurons expressed both *Trh* and *Adcyap1*, and most of the remaining *Sim2*+ neurons either expressed *Adcyap1* only (13.3 ± 1.2%) or *Trh* only (3.9 ± 1.1%) (Figure 1D). Interestingly, adult *Sim2*-expressing neurons were primarily located in the anterior and mid-PVH, with very few cells in the posterior PVH (Figure 1E), consistent with previous studies of newborn mice^33^. This anterior localization is also consistent with monosynaptic rabies mapping of ARC^AgRP^ neuron afferents^26^; and contrasts with PVH satiety neurons expressing *Mc4r* and *Pdyn*^24,25^, which are located primarily from mid-to posterior PVH.

To gain specific genetic access to PVH^Sim2^ neurons, we generated recombinase driver mice expressing *Cre* and *FLPo*, respectively, under the control of the endogenous *Sim2* gene (*Sim2-2a-Cre*, Figures S2A and S2B; and *Sim2-2a-FLPo*, Figures S3A and S3B). We generated both *Cre* and *FLPo* mouse lines to permit flexibility in experimental design. Using recombinase-dependent tdTomato reporter mice, we found that these lines drove expression in a number of brain regions, including in the PVH and mammillary bodies, where *Sim2* expression has been previously described (Figures S2C and S3C)^34,35^. Using smFISH for *Sim2* in these reporter mice, we verified a high degree of overlap between tdTomato and *Sim2* expression in the PVH (Figures S2D and S3D). Consistent with most PVH neurons being glutamatergic, we found that almost all PVH^Sim2^ neurons expressed *VGlut2 (Slc17a6)* (Figure S2D). Importantly, PVH^Sim2^ neurons are almost exclusively centrally-projecting, and not neuroendocrine, since the vast majority (88.6 ± 3.3%) of these neurons were not labelled by peripheral injection of fluorogold (Figure S4A; mean ± SEM, N = 4).

### PVH^Sim2^ neurons excite ARC^AgRP^ neurons and drive feeding

To test whether PVH^Sim2^ neurons send axonal projections to the mediobasal ARC, where ARC^AgRP^ neurons are located, we labelled PVH^Sim2^ neuron projections and ARC^AgRP^ neurons in the same mouse, using *Sim2-2a-Cre*::*Npy-hrGFP* mice (since ARC^AgRP^ neurons coexpress Npy^36^) injected with AAV-DIO-ChR2-mCherry in the PVH (Figure 1F). We then imaged a 700µm-thick section of ARC cleared using CUBIC^37^. We observed dense synaptic projections in the ARC, specifically in the region of ARC containing GFP+ ARC^AgRP^ neurons. Moreover, many GFP+ neurons in the ARC were densely wrapped by mCherry+ fibers, suggesting that PVH^Sim2^ neurons may provide direct excitatory input to ARC^AgRP^ neurons.

To more comprehensively visualize the synaptic projections of PVH^Sim2^ neurons, we injected an AAV expressing Cre-dependent, mCherry-tagged Synaptophysin (AAV-DIO-Syp-mCherry) into the PVH of *Sim2-2a-Cre* mice and examined labelled axonal projections throughout the brain (Figure S4B). This alternative approach confirmed that PVH^Sim2^ neurons are more prevalent in the anterior PVH, and that PVH^Sim2^ neuron projections densely and specifically target the mediobasal ARC. We also observed dense PVH^Sim2^ projections in the bed nucleus of the stria terminalis (BNST), as well as more sparse projections in a small number of other forebrain regions (Figure S4B).

Given that PVH^Sim2^ neurons as a whole project to multiple areas, individual PVH^Sim2^ neurons might project to individual target areas, or might collateralize to multiple downstream targets. Favoring the latter hypothesis, retrograde viral collateral mapping using AAVretro-fDIO-Cre revealed that at least a subset of PVH^Sim2^ neurons send collaterals to both the ARC and BNST (Figures S4C and S4D). This indicates at least some individual PVH^Sim2^ neurons project to multiple target sites, at least for the two densest projection targets.

The anatomy of PVH^Sim2^ neuron projections suggests that they are poised to form synaptic contacts with ARC^AgRP^ neurons. To test whether PVH^Sim2^ neurons provide direct monosynaptic input to ARC^AgRP^ neurons, we used channelrhodopsin (ChR2)-assisted circuit mapping (CRACM)^38^, using *Npy-hrGFP* to label ARC^AgRP^ neurons as above. In *ex vivo* brain slices, activating ChR2-expressing PVH^Sim2^ axons resulted in light-evoked excitatory post-synaptic currents (leEPSCs) with monosynaptic latency (∼6ms) in ∼40% of Npy+ neurons (Figure 1G). These leEPSCs were abolished by tetrodotoxin (TTX), but were reinstated by co-application of 4-aminopyridine (4-AP), confirming their monosynaptic nature (Figure 1H). No leEPSCs were observed in Npy-negative ARC neurons (Figure 1G). Of note, the observed PVH^Sim2^→ARC^AgRP^ connectivity rate is less than the ∼75% of Npy+ neurons that receive input from PVH^Trh^ neurons^28^, which may be due to lower ChR2 expression in *Sim2-2a-Cre* mice (since *Sim2* is expressed at lower levels than *Trh*, see expression levels in Figure 1B); or alternatively could suggest that PVH afferents to some ARC^AgRP^ neurons express *Trh* but not *Sim2*. Regardless, these results clearly establish that PVH^Sim2^ neurons provide selective monosynaptic glutamatergic input to ARC^AgRP^ neurons. Notably, given that the majority of PVH^Sim2^ neurons express *Adcyap1* (Figure 1D), and that PACAP (the neuropeptide product of *Adcyap1*) increases AgRP neuron firing rates *ex vivo*^26^, it is likely that PVH^Sim2^ neurons also influence AgRP neuron activity via neuropeptide release in addition to fast glutamatergic transmission.

To test whether PVH^Sim2^ neurons influence ARC^AgRP^ neuron activity *in vivo*, we optogenetically activated their projections to ARC while monitoring the activity of ARC^AgRP^ neurons using fiber photometry (Figure 1I). We expressed the red-shifted light-gated cation channel ChrimsonR in the PVH under the control of *Sim2-2a-Cre*, and the calcium-dependent fluorophore GCaMP6s in the ARC under the control of *Npy-IRES-FLPo* (since ARC^AgRP^ neurons coexpress Npy^36^), and then placed an optical fiber for light delivery and collection above the ARC. Brief (10s) stimulations of PVH^Sim2^ neurons increased ARC^AgRP^ neuron activity, which returned to baseline after stimulation offset (Figure 1I). Notably, this activity gradually increased over several seconds during the stimulation period. Thus, PVH^Sim2^ neurons can increase ARC^AgRP^ neuron activity *in vivo*. Of note, ARC^AgRP^ neurons project reciprocally to the PVH, where they inhibit satiety neurons and increase the activity of neurons controlling the hypothalamic-pituitary-adrenal axis^5,22,24,25^. PVH^Sim2^ neurons, however, are notably unaffected by ARC^AgRP^ neuron activation (Figures S5A and S5B), underlining the unidirectional nature of the PVH^Sim2^→ARC^AgRP^ neuron circuit.

Activation of ARC^AgRP^ neurons, or their PVH afferents labelled by *Trh-IRES-Cre* or *Pacap-IRES-Cre*, drives food intake^3,4,26^. Thus, we hypothesized that activating PVH^Sim2^ neurons would similarly increase food intake through activation of ARC^AgRP^ neurons. To test this, we expressed the excitatory chemogenetic receptor hM3Dq in PVH^Sim2^ neurons, and measured food intake in response to injection of vehicle or the hM3Dq ligand clozapine N-oxide (CNO) in the early light phase, when mice usually eat little (Figures 1J and 1K). Activation of PVH^Sim2^ neurons by CNO injection strongly increased food intake over 3 hours (Figure 1K). Next, to isolate the influence of ARC-projecting PVH^Sim2^ neurons, we specifically activated the PVH^Sim2^ neurons that project to the ARC. To achieve this, we injected AAVretro-fDIO-Cre into the ARC of *Sim2-2a-FLPo* mice to express Cre recombinase in ARC-projecting *Sim2*+ neurons, and then injected AAV expressing Cre-dependent, mCherry-tagged hM3Dq into the PVH (Figures S5C and S5D). Like activation of all PVH^Sim2^ neurons, activating just the ARC-projecting PVH^Sim2^ neurons by CNO injection increased food intake (Figure S5E). This indicates that PVH^Sim2^ neurons can drive food intake through the activation of ARC^AgRP^ neurons (Figure 1L). In support of this, inhibition of ARC^AgRP^ neurons has been shown to prevent the ability of PVH afferents (PVH^Trh^ neurons) to drive food intake^26^.

As noted above (Figure 1C), a small proportion of *Trh+/Adcyap1+* neurons do not express *Sim2*. Our sc/snRNA-seq atlas suggested that a small subset of *Trh*/*Adcyap1*-coexpressing PVH neurons belong to a cluster uniquely marked by expression of *Ucn3*, consistent with previous reports describing expression of *Trh* and *Adcyap1* in *Ucn3*+ neurons^30,39,40^. However, PVH^Ucn3^ neurons project weakly to the ARC, do not synapse onto ARC^AgRP^ neurons, and do not increase food intake when activated *in vivo* (Figure S6). Ucn3 expression therefore does not label the excitatory PVH input to ARC^AgRP^ neurons.

### Pre-ingestive inhibition of PVH^Sim2^ neurons by food

Having identified a specific marker for the PVH neurons that provide excitatory afferent input to ARC^AgRP^ neurons (Sim2) and generated recombinase driver mouse lines to provide highly selective access to them (*Sim2-2a-Cre* and *Sim2-2a-FLPo*), we can now study the information that these inputs convey to ARC^AgRP^ neurons by recording their activity *in vivo*. We started by characterizing the response of PVH^Sim2^ neurons to food, given that ARC^AgRP^ neurons show a strong reduction in activity in response to food in the fasted state^7–9^. To do so, we expressed GCaMP6s in PVH^Sim2^ neurons and monitored population calcium activity using fiber photometry (Figures 2A and 2B). Mice were fasted overnight, and then presented with a pellet of standard chow (∼2 g; Figure 2C).

**Figure 2.**
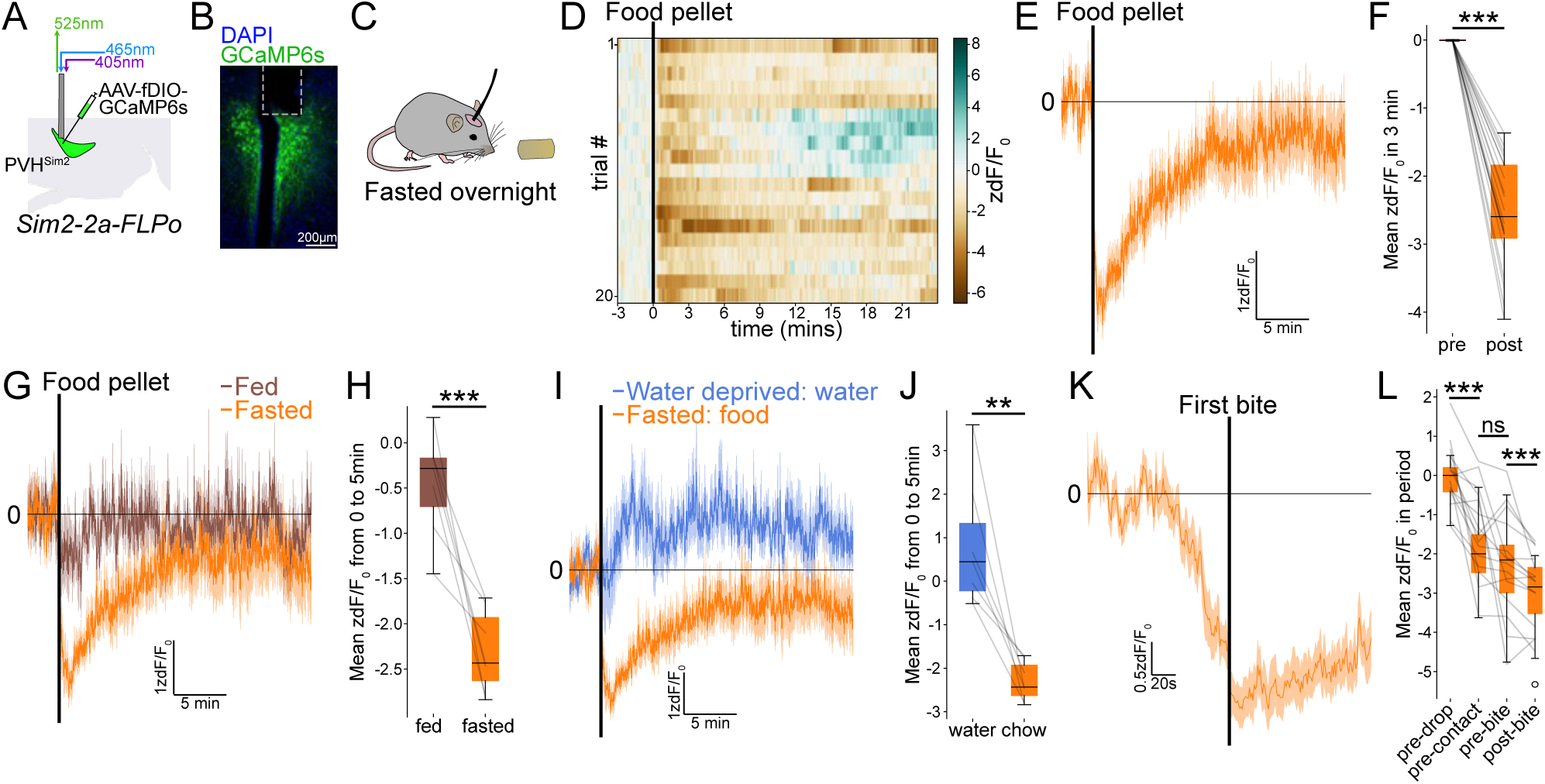
Pre-ingestive inhibition of PVH^Sim2^ neurons by food. A) Schematic of fiber photometry recordings from PVH^Sim2^ neurons. B) Representative image of GCaMP6s expression (green) in PVH^Sim2^ neurons; dashed line indicates optic fiber placement. C) Schematic of experimental setup: a fasted mouse is presented with a chow pellet. D) Heatmap of PVH^Sim2^ neuron activity before and after presentation of a chow pellet. Each row is one trial; black line indicates time of pellet delivery. (N=20 trials from 7 mice). E) Mean (± SEM) response of PVH^Sim2^ neurons to pellet delivery. F) Mean activity in 3 minutes before and after pellet delivery. Each line indicates one trial; box and whiskers represent upper/lower quartiles and minimum/maximum data points, respectively. (N = 20 trials from 7 mice; two-tailed paired t-test). G, I) Response of PVH^Sim2^ neurons to pellet delivery in the fed or fasted state (G), or to water bowl delivery in the water-deprived state and pellet delivery in the fasted state (I). (N = 7 mice; mean ± SEM over mice). H, J) Mean activity in 5 minutes following pellet delivery in fed and fasted states (H), or following water bowl delivery in the water-deprived state and pellet delivery in the fasted state (J). Each data point represents mean over trials for one mouse; lines connect data points for the same mouse. (N=7 mice; two-tailed paired t-test). K) Mean ± SEM activity of PVH^Sim2^ neurons as in (E), but aligned to the first bite of chow. L) Mean activity in different bins relative to pellet delivery, contact, and consumption (see Methods); each line indicates one trial. (N = 20 trials from 7 mice; two-way repeated measures ANOVA; post-hoc comparisons between adjacent time points with Sidak correction). **p<0.01; ***p<0.001; ns = not significant.

Presentation of food to a fasted mouse led to a rapid drop in PVH^Sim2^ activity (Figure 2D-F), which was complete within seconds (τ = 7.5 ± 6.2 s from pellet presentation, where τ represents the time at which 63.8% of the total fall in activity has occurred). This rapid drop in activity is reminiscent of the fall in ARC^AgRP^ neuron activity seen with food presentation^7–9^. Notably, PVH^Sim2^ activity gradually recovered following food presentation. This contrasts with the drop in ARC^AgRP^ neuron activity in response to food, which is sustained over at least tens of minutes after food consumption^7^. Thus, the initial fall in ARC^AgRP^ neuron activity is likely due, in part, to a reduction in excitatory input from PVH^Sim2^ neurons, in addition to an increase in inhibitory input from DMH^Lepr/Glp1r^ and ARC^Bnc2^ neurons^41–44^. The sustained reduction in ARC^AgRP^ neuron activity, however, is likely mediated by other inputs, such as post-ingestive signals originating in the gastrointestinal tract^10,11,45,46^.

As with ARC^AgRP^ neurons^7^, the food response of PVH^Sim2^ neurons was state-specific, since these neurons showed little response to food in the fed state (Figures 2G and 2H). Further, this response was specific to food, and not other motivationally salient stimuli, since following overnight water deprivation, presentation of water did not inhibit PVH^Sim2^ neuron activity (Figures 2I and 2J). In fact, water presentation slightly increased PVH^Sim2^ neuron activity, which may reflect increased food-seeking drive following the relief of dehydration-induced anorexia^47^.

The rapid response of PVH^Sim2^ neurons to food suggests that these neurons may be sensitive to sensory stimuli that are predictive of food. To further investigate this hypothesis, in a subset of mice, we recorded video together with fiber photometry, and manually annotated the interactions of mice with food. PVH^Sim2^ neuron activity dropped prior to the first bite of food, and even before the mouse made contact with the food pellet (Figures 2K and 2L). This indicates that, similar to ARC^AgRP^ neurons^7–9^, PVH^Sim2^ neurons are sensitive to pre-ingestive sensory cues that predict upcoming food intake.

### Removal of food rapidly increases ARC^AgRP^ and PVH^Sim2^ neuron activity

ARC^AgRP^ neurons are bidirectionally sensitive to energy status: while food presentation inhibits their activity, overnight fasting strongly increases the activity of ARC^AgRP^ neurons^8,48^. Using immediate early gene expression (Fos) as a readout of neuronal activity, we found that PVH^Sim2^ neurons similarly showed increased activity following overnight fasting (Figure S7). Thus, PVH^Sim2^ neurons likely contribute to the fasting activation of ARC^AgRP^ neurons.

We next sought to determine the timescale over which fasting activates ARC^AgRP^ neurons. Traditionally, the activation of ARC^AgRP^ neurons is thought to be driven largely by feedback from the body as energy stores are gradually depleted^1,49,50^. However, the PVH, where the Sim2+ inputs to ARC^AgRP^ neurons are located, is not thought to directly sense bodily feedback signals related to the fasted state, such as leptin^51^. Moreover, the timescale over which fasting activates ARC^AgRP^ neurons has remained unclear due to the inability to measure ARC^AgRP^ neuron activity continuously over hours during fasting. We recently developed a novel long-term fiber photometry approach to continuously measure population activity over days^17^. Using this methodology, we found that ARC^AgRP^ neurons show a remarkably rapid increase in activity when fasting is initiated by the removal of food^17^. This suggests that in addition to slower feedback signals from the body, rapid feedforward neuronal inputs play a role in the response of ARC^AgRP^ neurons to fasting.

To confirm these short– and long-term responses of ARC^AgRP^ neurons to fasting, we performed long-term fiber photometry recordings from ARC^AgRP^ neurons before, during, and after an overnight fast, as well as under *ad libitum* conditions (Figures 3A-D). We recorded activity from ZT6 (Zeitgeber Time 6; i.e., 6 hours after lights-on) until ZT5 the following day, and food was removed at ZT12 (lights-out) and returned at ZT2 (2 hours after lights-on). As previously described^17,52^, in the control condition with *ad libitum* access to food, ARC^AgRP^ neuron activity gradually decreased over the dark period (Figure 3E). In the overnight fast condition, however, removal of food led to a large, rapid, and sustained rise in ARC^AgRP^ neuron activity, within less than 1 hour of fasting onset (Figures 3E and 3F). This increase in activity was complete within ∼2 hours, with a time constant (τ) of 1.78 ± 0.40 h (mean ± SEM). The response of ARC^AgRP^ neurons happens before the body reaches a state of negative energy balance, since while 1 hour of fasting from lights-out did significantly increase ARC^AgRP^ neuron activity, it did not significantly affect circulating levels of ghrelin, insulin, or leptin (Figure S8). Thus, the activity of ARC^AgRP^ neurons *anticipates* the metabolic consequences of food unavailability early in fasting.

**Figure 3.**
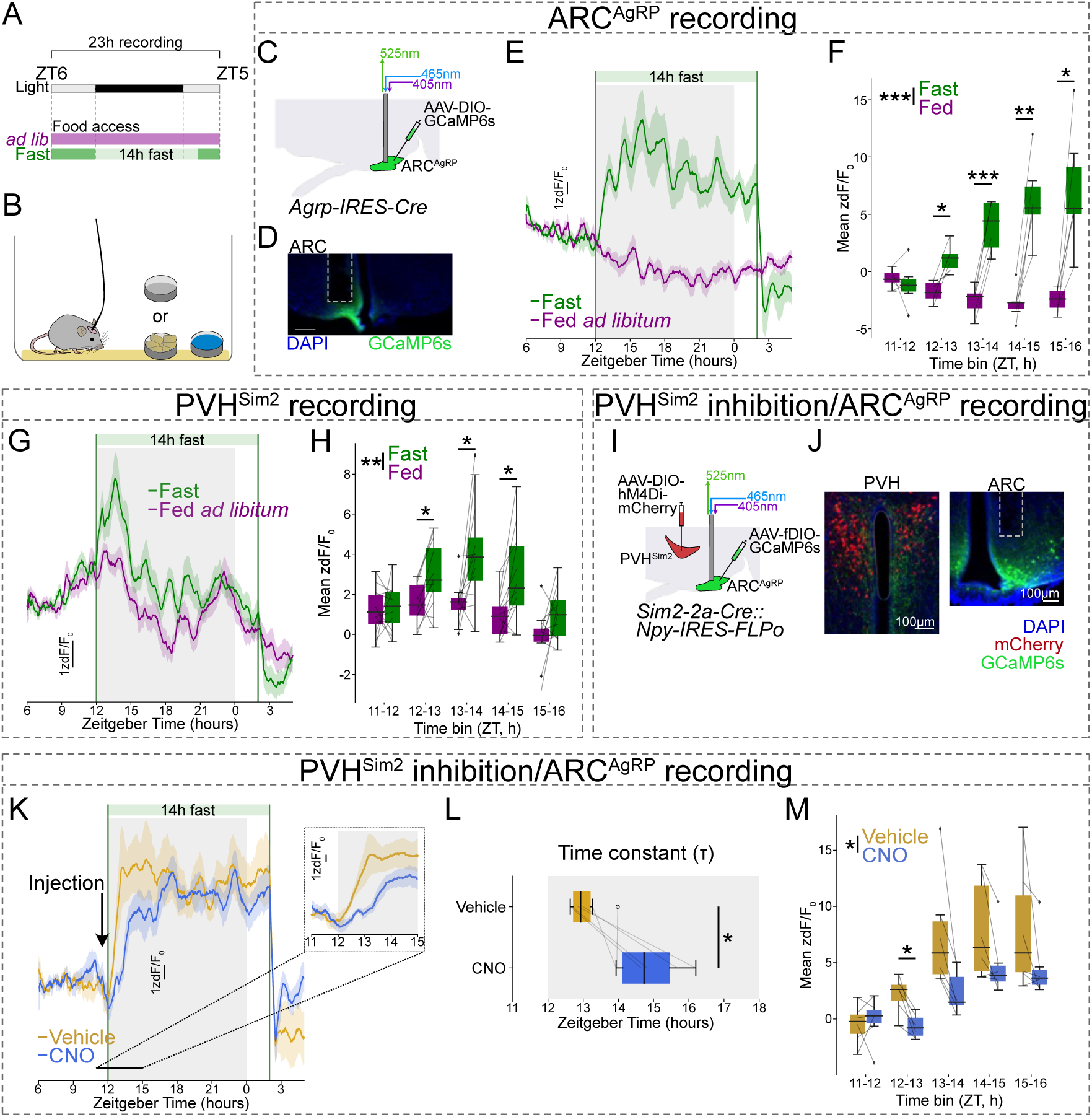
PVH^Sim2^ neurons mediate the rapid activation of ARC^AgRP^ neurons upon food removal. A) Schematic of experimental design: activity is recorded from ZT6 to ZT5 the following day (23 hour long recording), and in the fast condition, food is removed between ZT12 and ZT2 (14 hour fast). B) Experimental setup for home-cage recording. C) Schema for fiber photometry recording from ARC^AgRP^ neurons. D) Representative image of GCaMP6s expression (green) in ARC^AgRP^ neurons; dashed line indicates optic fiber placement. E) Recording of ARC^AgRP^ neuron activity in the fast and *ad libitum* conditions (mean ± SEM over mice; 30-minute moving mean). Vertical lines indicate the start and end of fasting. (N = 6 mice). F) Mean activity in 1-hour bins between ZT11 and ZT16. Two-way repeated measures ANOVA; post-hoc comparisons at each time point with Holm-Sidak correction. G, H) As (E, F) but for PVH^Sim2^ neurons in *Sim2-2a-FLPo* mice. (N = 12 mice). I) Schema for fiber photometry recording from ARC^AgRP^ neurons (using *Npy-IRES-FLPo*) with chemogenetic inhibition of PVH^Sim2^ neurons. J) Representative images of mCherry (red) and GCaMP6s (green) expression in the PVH (left) and ARC (right). K) Recording of ARC^AgRP^ neuron activity (mean ± SEM) in the fast condition with injection of vehicle or CNO at ZT11.5. Inset: magnification of activity between ZT11 and ZT15. (N = 6 mice). L) Time constant (τ) of increase in ARC^AgRP^ neuron activity for recordings shown in (K). Two-tailed paired t-test. M) As (F), but for the ARC^AgRP^ recordings with PVH^Sim2^ neuron inhibition presented in (K). Two-way repeated measures ANOVA; post-hoc comparisons at each time point with Holm-Sidak correction. *p<0.05; **p<0.01; ***p<0.001. For boxplots, lines indicate individual mice; box and whiskers indicate upper/lower quartiles and minimum/maximum data points, respectively. Outliers indicated by symbol.

Given that PVH^Sim2^ neurons provide excitatory input to ARC^AgRP^ neurons, we speculated that they may convey this rapid fasting response to ARC^AgRP^ neurons. To test whether PVH^Sim2^ neurons are similarly sensitive to food removal, we recorded PVH^Sim2^ neuron activity under the same conditions (Figures 3G and 3H). In the *ad libitum* state, PVH^Sim2^ activity was dynamic, peaking in the early dark phase. Strikingly, following food removal at ZT12, PVH^Sim2^ neuron activity increased rapidly (within 1 hour; τ = 0.87 ± 0.14 h (mean ± SEM)). Activity remained elevated above the *ad libitum* state for at least 3 hours following food removal, before then dropping. Consistent with the results presented in Figure 2, returning food at ZT2 led to a rapid drop in PVH^Sim2^ neuron activity. The rapid deprivation response is food-specific because similarly timed water deprivation did not increase ARC^AgRP^ or PVH^Sim2^ neuron activity relative to the *ad libitum* condition (Figures S9A-D). This indicates that food removal transiently increases PVH^Sim2^ neuron activity over a period of hours, and suggests that these neurons contribute to the fasting activation of ARC^AgRP^ neurons.

In addition to excitatory input from the PVH, ARC^AgRP^ neurons also receive strong inhibitory drive from *Lepr/Glp1r*-expressing neurons in the ventral dorsomedial hypothalamus (vDMH^LepR^)^41,44^, and the rapid response of these neurons to food presentation contributes to the inhibition of ARC^AgRP^ neurons by feedforward cues that predict food^42^. To test whether vDMH^LepR^ neurons might also contribute to the increase in ARC^AgRP^ neuron activity early in fasting, we monitored their activity under the same conditions (Figures S9E and S9F). As previously reported^41^, vDMH^LepR^ activity strongly increased when food was presented at the end of fasting (Figure S9G). However, we observed no difference in vDMH^LepR^ activity between *ad libitum* and fasted states within the first 6 hours of fasting (Figures S9G and S9H), indicating that vDMH^LepR^ neurons do not contribute to the activation of ARC^AgRP^ neurons by food removal.

### PVH^Sim2^ neurons mediate the rapid ARC^AgRP^ neuron response to food removal

Given that the activity of both PVH^Sim2^ and ARC^AgRP^ neurons increases in response to food removal, and that PVH^Sim2^ neurons project to and activate ARC^AgRP^ neurons, we hypothesized that PVH^Sim2^ neurons drive the ARC^AgRP^ neuron response to food removal. To test this, we inhibited PVH^Sim2^ neuron activity while recording ARC^AgRP^ neuron activity (Figures 3I and 3J). In *Sim2-2a-Cre*::*Npy-IRES-FLPo* mice, we expressed the inhibitory chemogenetic receptor hM4Di in PVH^Sim2^ neurons using a Cre-dependent AAV. We then used a FLP-dependent AAV to express GCaMP6s in ARC^AgRP^ neurons, and implanted an optical fiber above the ARC for fiber photometry recordings. We recorded ARC^AgRP^ neuron activity before, during, and after an overnight fast as described above in Figure 3A, and injected clozapine N-oxide (CNO) or vehicle at ZT11.5 – i.e., 30 minutes before food removal. Following vehicle injection, ARC^AgRP^ neurons responded to food removal with a rapid, large increase in activity (Figures 3K-M). Inhibition of PVH^Sim2^ neurons by injection of CNO delayed this increase in activity, increasing the time constant (τ) by an average of 1.80 ± 0.51 hours (mean ± SEM). Likewise, inhibiting PVH inputs to ARC^AgRP^ neurons using the less-specific but broader *Trh-IRES-Cre* mouse line similarly delayed the increase in ARC^AgRP^ neuron activity early in fasting (Figures S10A-E; Δτ = 2.03 ± 0.34 h (mean ± SEM)). Thus, the rapid increase in ARC^AgRP^ neuron activity early in fasting is driven by excitatory input from PVH^Sim2^ neurons.

ARC^AgRP^ neuron activity is essential for the high level of food intake displayed following an overnight fast^53^. To test whether PVH^Sim2^ neurons are similarly required for post-fast feeding, we acutely inhibited PVH^Sim2^ neurons the morning after an overnight fast.

Chemogenetic inhibition of PVH^Sim2^ neurons did not affect food intake in the first 3 hours of refeeding (Figures S10F and S10G). This suggests that other factors (e.g. circulating hormones) may sustain ARC^AgRP^ neuron activity and drive food intake following long-term fasting. Notably, acute inhibition of PVH^Trh^ neurons does reduce feeding early in the dark cycle^26^, suggesting that PVH afferents contribute to the control of *ad libitum* food intake (see also Figure 5, below).

In the experiments presented above, fasting was always initiated at ZT12, which is when mice usually begin to eat at the highest rate^17^. We wondered whether the neuronal response to food removal might change if we removed food at different times relative to the light cycle. To test this, we removed food either 3 or 6 hours before the dark phase (ZT6 or ZT9, respectively), and recorded ARC^AgRP^ and PVH^Sim2^ neuron activity (Figures S11A-C). Food removal at ZT9 did lead to small increases in ARC^AgRP^ and PVH^Sim2^ neuron activity within the subsequent 3 hours, relative to the *ad libitum* state. However, no matter when food was removed, both ARC^AgRP^ and PVH^Sim2^ neuron activity in the fasted state always peaked early in the dark phase (Figures S11D and S11E), with ARC^AgRP^ neuron activity peaking around 2 hours later than that of PVH^Sim2^ neurons. The consistency in the timing of peak ARC^AgRP^ and PVH^Sim2^ neuron activity irrespective of fast start time suggests that it is not the removal of food *per se* that most potently increases the activity of these neurons. Rather, the early dark phase is when mice usually eat at the highest rate, and thus have the highest motivational drive to find and consume food. Thus, this suggests that the unavailability of food *at the time when mice have a strong drive to eat* triggers the rapid increase in PVH^Sim2^ and ARC^AgRP^ neuron activity.

### Unsuccessful food-seeking increases PVH^Sim2^ and ARC^AgRP^ neuron activity

The results presented above suggest that increased PVH^Sim2^ and ARC^AgRP^ neuron activity early in fasting may be triggered by the unavailability of food at times when mice desire food and are therefore engaged in food-seeking behavior. In other words, ARC^AgRP^ neuron activity may increase as mice realize that food is unavailable, because their search for food is unsuccessful. In our prior experiments, however, we were unable to know when food seeking occurred and was unsuccessful. To establish when mice realize that food is unavailable, we measured food-seeking behavior during fasting. To do so, we trained mice to engage in nose-poking behavior in order to obtain food in their home cage, using an operant feeding device (FED3)^54^. Mice were trained to receive all their food by performing an operant task on a fixed-ratio 1 (FR1) schedule, where each poke leads to delivery of one small (20 mg) food pellet. Then, we recorded ARC^AgRP^ neuron activity in mice either with constant access to food on a FR1 schedule or throughout an overnight fast, during which mice could still poke, but no food pellets were delivered (Figures 4A and 4B).

**Figure 4.**
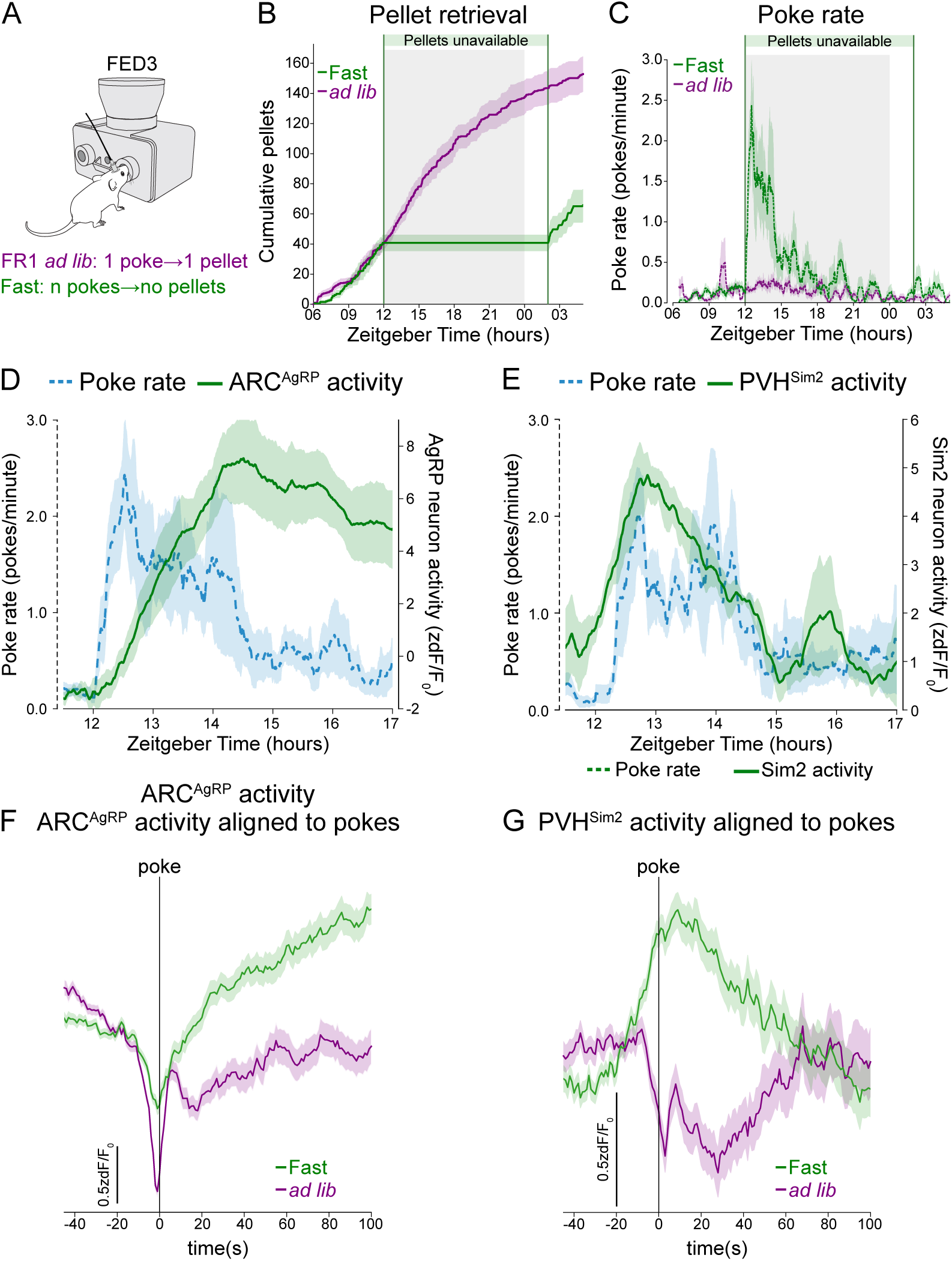
Unsuccessful food seeking increases ARC^AgRP^ and PVH^Sim2^ neuron activity. A) Schema of operant task using FED3: in the *ad libitum* state, each nose poke leads to the delivery of one food pellet. During fasting (ZT12-ZT2, 14 hour fast), mice can still poke, but no pellets are delivered. B) Cumulative number of pellets retrieved in fast (green line) and *ad libitum* (purple line) conditions (mean ± SEM). C) Rolling mean of poke rate in fast (dotted green line) and *ad libitum* (dotted purple line) conditions (mean ± SEM). D) Overlay of poke rate (dotted blue line) and ARC^AgRP^ neuron activity (solid green line) (mean ± SEM over mice; 30-minute moving mean) during the first 5 hours of fasting. E) Overlay of poke rate (dotted blue line) and PVH^Sim2^ neuron activity (solid green line) during the first 5 hours of fasting. B-E: mean ± SEM over mice. F,G) ARC^AgRP^ (F) and PVH^Sim2^ (G) neuron activity aligned to pokes (mean ± SEM over pokes). B,C,D,F: N = 7 mice; E,G: N = 4 mice.

**Figure 5.**
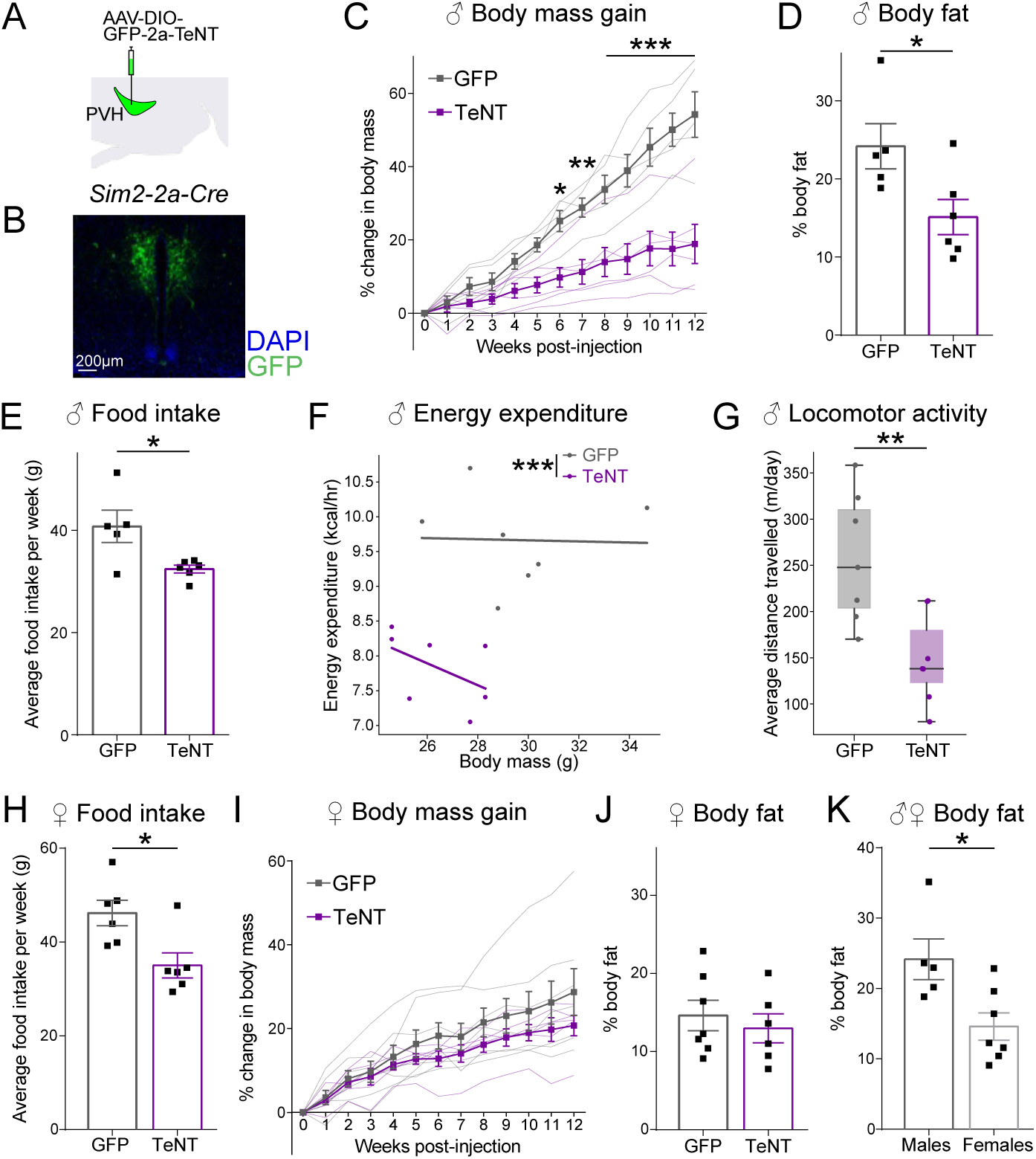
PVH^Sim2^ neurons are necessary for normal food intake. A) Schematic of AAV-DIO-GFP-2a-TeNT injection into the PVH of *Sim2-2a-Cre* mice. B) Representative image of resultant GFP expression (green) in the PVH. C) Percentage change in body mass of males following AAV injection, relative to pre-injection. Mean ± SEM; thin lines represent individual mice. (N = 5-6 mice per condition; two-way repeated measures ANOVA; post-hoc comparisons at each time point with Sidak correction). D) Fat mass as a percentage of total body mass in male mice at 13 weeks post-injection. (Mean ± SEM, N = 5-6 mice per condition; two-tailed unpaired t-test). E) Mean weekly food intake of male mice from weeks 4-12 post-injection. (Mean ± SEM, N = 5-6 mice per condition; two-tailed unpaired t-test). F) Mean hourly energy expenditure (over two days) vs body mass of males at 4 weeks post-injection for each mouse. Lines represent linear regression fit for each group. (N = 7 mice per condition; generalized linear model). G) Mean distance travelled per day (over two days) at 4 weeks post-injection. Box/whiskers represent upper/lower quartiles and minimum/maximum values, respectively. (N = 7; two-tailed unpaired t-test). H) Mean weekly food intake of female mice from weeks 4-12 post-injection. (Mean ± SEM, N = 6-7 mice per condition; two-tailed unpaired t-test). I) Percentage change in body mass of females following AAV injection, relative to pre-injection. Mean ± SEM; thin lines represent individual mice. (N = 6-7 mice per condition; two-way repeated measures ANOVA: no significant effect of AAV or interaction). J) Fat mass as a percentage of total body mass in female mice at 13 weeks post-injection. (Mean ± SEM, N = 6-7 mice per condition; two-tailed unpaired t-test). K) Fat mass as a percentage of total body mass in GFP-injected control male and female mice at 13 weeks post-injection. (Mean± SEM, N = 5-7 mice per condition; two-tailed unpaired t-test).

At the beginning of fasting, mice engaged in vigorous poking at a high rate, which then gradually declined over hours (Figure 4C). This gradual decline in poking represents extinction, the gradual weakening of operant nose-poking behavior as mice learn that this behavior no longer results in food delivery. The initial high poke rate, meanwhile, is an extinction burst: an initial increase in operant behavior that is frequently observed early in extinction^55^. This extinction burst is a behavioral indication that the mice have noticed the change in contingency, while the decline in poke rate shows that they are learning the new contingency (i.e. that pokes no longer deliver food). Similar to the fasted condition in our previous experiments, in the operant context ARC^AgRP^ neuron activity increased early in fasting, and then remained elevated (Figure 4D). Notably, this increase in ARC^AgRP^ neuron activity coincided with the decline in the rate of food-seeking (nose-poking) behavior (Figure 4D). As such, ARC^AgRP^ neuron activity increases as mice enter a state of awareness that food is not available, and we speculate that this awareness drives the increase in ARC^AgRP^ neuron activity.

Given that PVH^Sim2^ neurons are also active early in fasting, we reasoned that they may similarly respond to unsuccessful food-seeking. Like ARC^AgRP^ neurons, the activity of PVH^Sim2^ neurons was also elevated during the initial burst of poking early in fasting, demonstrating that PVH^Sim2^ neurons are also active during unsuccessful food-seeking behavior (Figure 4E). However, ARC^AgRP^ neurons and PVH^Sim2^ neurons showed different temporal dynamics early in fasting: while ARC^AgRP^ neuron activity increased gradually as poke rate declined, and then remained elevated, PVH^Sim2^ neuron activity increased more rapidly and was high throughout the period of high poke rate, but then declined together with the poke rate. This indicates that PVH^Sim2^ neurons respond transiently to food unavailability, and suggests that ARC^AgRP^ neurons may use these transient signals to obtain a stable prediction of the availability of food.

To gain a closer understanding of how the awareness of food unavailability impacts ARC^AgRP^ and PVH^Sim2^ neuron activity, we looked at the response of these neuronal populations to individual pokes over a shorter timescale. We aligned neural activity to pokes, selecting pokes only during the fasting or equivalent *ad libitum* period (ZT12-ZT2). To avoid including preceding pokes during the baseline period, we selected only those pokes that were not preceded by another poke within 45 s, although this filter did not affect the results of this analysis (Figure S12). Under *ad libitum* conditions, ARC^AgRP^ neuron activity dropped precipitously immediately preceding a poke, and then recovered to a level below its initial baseline (Figure 4F). This is consistent with the reduction in ARC^AgRP^ neuron activity observed prior to feeding bouts in mice with free access to food^17^. During fasting, however, this initial drop was reduced in amplitude, perhaps because the expectation of a poke leading to food was reduced; and notably, the unrewarded poke was followed by a large increase in ARC^AgRP^ neuron activity. This response to unrewarded pokes was fairly consistent throughout an overnight fast (Figure S13A). Thus, unsuccessful food-seeking behavior increases ARC^AgRP^ neuron activity.

Like ARC^AgRP^ neurons, PVH^Sim2^ neuron activity dropped at the time of poking when food was available (Figure 4G). In contrast, during fasting PVH^Sim2^ neuron activity increased with unrewarded pokes. This response occurred earlier than that in ARC^AgRP^ neurons; in fact, increased PVH^Sim2^ neuron activity preceded pokes during fasting. This increase in PVH^Sim2^ neuron activity before poking developed gradually over the course of fasting: early in fasting, PVH^Sim2^ neuron activity increased only after poking, once food was not delivered; but this response shifted to earlier time points as fasting progressed (Figure S13B). We speculate that later in fasting, PVH^Sim2^ neurons show an anticipatory response that reflects the prediction that food will not be delivered. Thus, both PVH^Sim2^ and ARC^AgRP^ neurons show increased activity in response to unsuccessful food seeking, with PVH^Sim2^ neurons showing more rapid responses, particularly early in fasting.

### PVH^Sim2^ neurons are necessary for normal food intake

We demonstrated above that PVH^Sim2^ neuron activity is bidirectionally sensitive to the availability of food, rapidly decreasing when food is unexpectedly present (Figure 2) and rapidly increasing when it is unexpectedly absent (Figures 3 and 4). We next asked how the ongoing activity of PVH^Sim2^ neurons contributes to long-term energy balance in mice with *ad libitum* access to food. To do so, we chronically silenced synaptic output from PVH^Sim2^ neurons by injecting AAV-DIO-GFP-2a-TeNT into the PVH of *Sim2-2a-Cre* mice (Figures 5A and 5B). We then measured body weight and food intake weekly over 12 weeks, relative to control (GFP-injected) mice. While control males gained weight throughout the experiment, silencing PVH^Sim2^ output drastically reduced body weight gain (Figure 5C). Moreover, at the end of this experiment, PVH^Sim2^-TeNT males had reduced body fat relative to controls (Figure 5D). Consistent with the ability of these neurons to drive food intake when activated (Figure 1K), we found that chronic silencing of PVH^Sim2^ output reduced food intake (Figure 5E, average of weeks 4-12 post-injection).

To test whether the reduced body weight of PVH^Sim2^-TeNT male mice might be due to this reduction in food intake, or to metabolic alterations, we used indirect calorimetry to measure a variety of metabolic parameters at 4 weeks post-injection (when the substantial difference in weight is beginning to develop, Figure S14A). Silencing PVH^Sim2^ output greatly reduced energy expenditure (more than expected based on body weight, Figures 5F and S14B), as well as locomotor activity (Figures 5G and S14C), without affecting substrate utilization (as measured by respiratory exchange ratio, Figure S14D). All else being equal, this reduced metabolic activity would be expected to increase weight gain – in contrast to the reduced weight gain observed in PVH^Sim2^-TeNT males.

This indicates that the reduced weight following PVH^Sim2^ silencing in male mice is due to reduced food intake. Of note, reduced energy expenditure is unlikely to be caused by reduced ARC^AgRP^ neuron activity, since this is expected to increase energy expenditure^4,56^. Instead, we propose that this reduction in energy expenditure may be a metabolic adaptation to minimize negative energy balance in the face of this reduced energy consumption.

We also measured the effect of PVH^Sim2^ synaptic silencing on body weight and food intake in female mice. As in males, chronic silencing of PVH^Sim2^ output greatly reduced food intake (Figure 5H). Unlike in males, however, this reduced food intake did not translate to a statistically significant reduction in body weight or body fat (Figures 5I and 5J). This is likely due to a concurrent reduction in energy expenditure (as seen in males, Figure 5F), which was perhaps more effective in maintaining body weight due to the greater compensatory responses of these female mice to defend their much lower body fat content (Figure 5K: GFP controls measured at 13 weeks post-injection). Taken together, PVH^Sim2^ neuron activity is essential for normal levels of food intake in *ad libitum*-fed mice.

## Discussion

When faced with a shortage of food, animals display a variety of physiological and behavioral adaptations, including increased food-seeking and, once food is available, increased food intake. ARC^AgRP^ neurons are a key player in this fasting response, but how their activity is regulated, particularly during fasting, is unclear. ARC^AgRP^ neurons are classically thought to be activated in response to bodily energy deficit by circulating factors in the blood^1,2^. However, ARC^AgRP^ neurons also receive direct neuronal input from a variety of brain regions. One of the primary sources of neuronal input to ARC^AgRP^ neurons comes from excitatory neurons in the PVH^26^; and indeed, these PVH→ARC^AgRP^ synaptic connections undergo profound plasticity during negative energy balance^27,28^.

However, the precise molecular identity of PVH inputs to ARC^AgRP^ neurons, the information that they convey, and their role in the regulation of ARC^AgRP^ neuron activity and food intake, are still unknown.

Here, we began by interrogating a single-cell/nucleus RNA-seq atlas to identify the PVH transcriptomic neuron subtype that provides this excitatory drive to ARC^AgRP^ neurons.

PVH neurons expressing *Sim2* synapse onto and excite ARC^AgRP^ neurons, and drive food intake. We show that PVH^Sim2^ neuron activity is bidirectionally regulated by the availability of food: presentation of food to a fasted mouse markedly decreases PVH^Sim2^ neuron activity over minutes (Figure 2), whereas removal of food at times when mice would usually eat sharply increases PVH^Sim2^ neuron activity over 30-60 minutes (Figure 3). This latter response is triggered by unsuccessful food seeking behavior (Figure 4), perhaps explaining why it is most prominently seen at times when mice are more likely to eat (Figure S11). Moreover, we demonstrate that PVH^Sim2^ neuron activity is required for the rapid, feedforward activation of ARC^AgRP^ neurons early in fasting. Lastly, we show that at longer timescales, PVH^Sim2^ neuron activity is required to support *ad libitum* food intake (Figure 5).

The neuron population identified in this study is transcriptomically, anatomically, and functionally unique among PVH neurons. While essentially all PVH neurons express the *Sim1* transcription factor^32^, both during development and adulthood, expression of its paralogue *Sim2* is highly specific to a subset of anterior PVH neurons co-expressing *Trh* and *Adcyap1* (Figures 1C-E). The anterior localization of PVH^Sim2^ neurons contrasts with other *Trh*-expressing PVH neuron populations^57^. Moreover, PVH^Sim2^ neurons largely project to targets within the hypothalamus and forebrain, with very few caudal projections (Figures 1 and S4). This contrasts with most previously described markers of subsets of PVH neurons, which project heavily to midbrain, brainstem, and spinal cord^23–25,58–64^. Interestingly, most of these other PVH populations also suppress food intake when activated (for example, those labelled by expression of *Mc4r*, *Pdyn*, *TrkB*, *Glp1r*, *Calcr*, *Irs4*, *Avp*, and *Nos1*)^23–25,58–60,63,65^, in contrast with PVH^Sim2^ neurons, which increase food intake, further highlighting how *Sim2*-expressing neurons play a unique role in energy balance regulation within the PVH.

Like the vast majority of PVH neurons, PVH^Sim2^ neurons are glutamatergic. In addition, PVH^Sim2^ neurons express multiple neuropeptide genes, including *Trh* and *Adcyap1* (encoding the neuropeptide PACAP, Figure 1D). This suggests that they may regulate ARC^AgRP^ neuron activity not only through glutamate release, but also through peptidergic transmission, particularly since PACAP can increase the firing rate of ARC^AgRP^ neurons^26^. Of note, the Sim2 transcription factor likely plays a role in the development of this PVH subpopulation, since knockout of *Sim2* reduces *Trh* expression in the anterior PVH^33^. The role of *Sim2* expression in adulthood in this neuron population, however, is unclear. Future studies will be necessary to determine whether *Sim2* expression in the adult PVH is required for the function of this population, and for energy balance regulation.

Predicting future energy states allows the brain to anticipate and prevent deviations from homeostasis before they occur^66^. In the context of hunger, a number of feedforward signals are known to regulate the activity of ARC^AgRP^ neurons in anticipation of future energy deficit or surfeit. First, following a period of fasting (during which ARC^AgRP^ neuron activity is elevated), presentation of food leads to a rapid and sustained drop in ARC^AgRP^ neuron activity, driven by external sensory cues predictive of food, the taste of food in the mouth, and post-ingestive detection of nutrients in the gastrointestinal tract^7–11,45,46,67^. Inhibitory inputs to ARC^AgRP^ neurons from both the DMH (DMH^Lepr/Glp1r^ neurons) and ARC (ARC^Bnc2^ neurons) show increased activity in response to sensory cues predictive of food^41–44^. Our results suggest that a drop in excitatory input to ARC^AgRP^ neurons likely also contributes to the food cue-induced inhibition of AgRP neurons, since the activity of PVH^Sim2^ neurons drops rapidly and markedly in response to food presentation before contact with food. Thus, a coordinated response involving an increase in inhibitory input with a concomitant decrease in excitatory input likely mediates the rapid feedforward inhibition of AgRP neurons. Interestingly the temporal dynamics of activity changes in the different inputs are not exactly alike: the response of PVH^Sim2^ neurons to food appears to be more sustained than that of DMH^Lepr/Glp1r^ neurons^41,42^, but less sustained than the prolonged food response of ARC^Bnc2^ neurons^43^.

A second component of the feedforward regulation of ARC^AgRP^ neuron activity is the rapid increase in their activity over the first hour of fasting (Figure 3 and 17), which occurs well before bodily energy stores are reduced. Here, we show that this increased activity is likely driven by unsuccessful food-seeking, which allows mice to become aware that food is unavailable (Figure 4). This response of ARC^AgRP^ neurons is driven by PVH^Sim2^ neurons, which are also rapidly activated by food unavailability, since inhibition of PVH^Sim2^ neuron activity delays the increase in ARC^AgRP^ neuron activity by around 2 hours. Of note, we cannot rule out a role for PVH^Sim2^ neurons in sustaining high ARC^AgRP^ neuron activity later in fasting, since a single CNO injection is unlikely to inhibit the activity of PVH^Sim2^ neurons throughout a 14-hour fast^68^.

What might be the functional role of the rapid PVH^Sim2^/ARC^AgRP^ neuron activation in response to food removal? ARC^AgRP^ neuron activation drives food-seeking and consumption^4^, so this rapid fasting response would be expected to drive mice to increase efforts to seek out food. Food-seeking in anticipation of a predicted energy deficit would potentially drive mice to find and consume food *before* they ever reach this energy deficit, and thus facilitate the maintenance of energy homeostasis. In addition, ARC^AgRP^ neurons also drive a number of physiological responses to fasting, including reduced energy expenditure, hypothalamic-pituitary-adrenal axis activation, peripheral insulin sensitivity, and autophagy in the liver^4,5,69,70^. The induction of these physiological responses in anticipation of energy deficit may help to conserve energy stores and maintain blood glucose as energy deficit develops during extended periods of fasting. Notably, such anticipatory responses to predict and prevent deviations from homeostasis are widespread across species^66^.

Importantly, increased activity in PVH inputs to ARC^AgRP^ neurons has been shown to drive synaptic plasticity, increasing the strength of PVH→ARC^AgRP^ connections by increasing the number of synaptic contacts from PVH^Trh^ to ARC^AgRP^ neurons^28^. Likewise, fasting similarly increases the strength of this connection, an effect that depends on NMDA receptor expression in ARC^AgRP^ neurons^27,28^. The substantial increase in PVH^Sim2^ neuron activity we observe early in the dark phase during fasting (Figure 3) raises the intriguing possibility that this high PVH^Sim2^ activity may be responsible for driving synaptic plasticity early in fasting. Of note, PVH^Sim2^ activity peaks earlier in fasting than ARC^AgRP^ neuron activity (Figure S11), consistent with the idea that strengthening of PVH^Sim2^→ARC^AgRP^ neuron connections may contribute to the early response of ARC^AgRP^ neurons to fasting. This strengthened PVH^Sim2^→ARC^AgRP^ neuron connection could plausibly then contribute to sustaining the high level of ARC^AgRP^ neuron activity throughout fasting, even as the activity of PVH^Sim2^ neurons drops. Notably, our sc/snRNA-seq data indicates that PVH^Sim2^ neurons express a number of mediators of excitatory synaptic plasticity at high levels, including *Bdnf* and *Cbln2*, a result that we confirmed with *in situ* hybridization (Figure S15). Indeed, within the PVH, *Cbln2* expression is largely selective to PVH^Sim2^ neurons. Future studies will be necessary to determine whether PVH^Sim2^ neuron activity early in fasting drives plasticity at PVH^Sim2^→ARC^AgRP^ neuron synapses, the molecular mechanisms responsible for this plasticity, and its role in sustaining ARC^AgRP^ neuron activity during fasting.

A third source of feedforward input to ARC^AgRP^ neurons is the circadian clock, which regulates ARC^AgRP^ neuron activity according to the time of day in order to appropriately time feeding behavior^17,52^. We similarly observed variability in PVH^Sim2^ neuron activity across the day under *ad libitum* conditions, with a peak in the early dark phase (Figure 3). However, PVH^Sim2^ neurons are unlikely to be responsible for circadian regulation of ARC^AgRP^ neuron activity, since a recent study found that ablation of PVH^Trh^ neurons did not affect ARC^AgRP^ neuron rhythmicity under *ad libitum* conditions^52^. Indeed, we recently demonstrated that this circadian rhythmicity in ARC^AgRP^ neuron activity is mediated by excitatory input from the DMH, which transmits circadian signals originating in the suprachiasmatic nucleus^17^. This is consistent with the idea that PVH^Sim2^ neurons are not required for the circadian regulation of ARC^AgRP^ neuron activity under *ad libitum* conditions, but instead play a specific role in the rapid response of ARC^AgRP^ neurons to fasting when food is desired but unavailable.

Beyond the feedforward regulation of hunger and satiety states, our results demonstrate that PVH^Sim2^ neurons also play a key role in regulating long-term levels of food intake under *ad libitum* conditions (Figure 5), since synaptic silencing of PVH^Sim2^ neuron output led to a marked reduction in food intake in both male and female mice. We propose that this effect of chronic loss of PVH^Sim2^ neuron output on food intake is due to a reduction in excitatory input to ARC^AgRP^ neurons. While chronic silencing or ablation of ARC^AgRP^ neurons in adult mice has been shown not to affect body weight gain or food intake under *ad libitum* conditions^53,71^, reduced excitatory input to ARC^AgRP^ neurons due to AgRP neuron-specific knockout of NMDA receptors does substantially reduce body weight and food intake^27^. This suggests that a reduction in excitatory input to ARC^AgRP^ neurons is likely to account for the reduced food intake of mice lacking PVH^Sim2^ neuron output.

In summary, we have identified a highly specific subtype of PVH neurons that drives food intake and is essential to support long-term energy balance. Further, we show that these PVH^Sim2^ neurons transmit bidirectional anticipatory signals to enable ARC^AgRP^ “hunger” neurons to predict future changes in energy balance, both when food is unexpectedly available, and when desired food is unavailable. Our findings demonstrate the importance of cognitively-processed, feedforward information in the control of hunger and energy balance, and provide an entry point to understand how the brain computes these feedforward signals. These results also identify a possible new neuronal target for cell-type-specific therapies that could potentially sustain reduced body weight after dieting.

## Acknowledgements

We thank all members of the B.B.L. laboratory for helpful discussions. We thank the BNORC transgenic core (NIH P30DK057521 and P30DK046200) for performing embryo injections to generate knockin mouse lines. We also thank the BIDMC Energy Balance Core (supported by NIH S10OD028635 and the Boston Area Diabetes Endocrinology Research Centers, P30DK135043), where Marissa Cortopassi performed indirect calorimetry experiments, and Alexander Banks assisted with data analysis and interpretation. Confocal imaging was performed at BIDMC’s Confocal Imaging Core. We thank Hakan Kucukdereli for assistance with optogenetic experiments, and Chen Wu for assistance in designing knock-in mouse lines. This work was supported by the NIH (R01DK134427, R01DK096010, and R01DK075632 to B.B.L.). Authors were supported by an EMBO Long-Term Fellowship (770-2018, S.J.W.), a T32 Postdoctoral Training Fellowship (5T32DK007516, E.D.L.), the Charles A. King Trust Postdoctoral Research Fellowship Program (A.M.D.), and a K99 Career Development Award (K99HL144923, J.M.R.).

## Author contributions

S.J.W. and B.B.L. conceived the study, designed the experiments, and wrote the manuscript with input from all authors. J.C.M. performed and analyzed all electrophysiological experiments. E.D.L. and J.T. performed *in situ* hybridization. J.M.R. led the PVH sc/snRNAseq experiments and analysis that formed the basis of this work.

S.J.W. and E.D.L. performed all other experiments and data analysis, with technical support from A.M.D.. S.J.W. and E.D.L. prepared figures.

## Supplemental information

Document S1. Figures S1-S15.

## STAR Methods

### RESOURCE AVAILABILITY

#### Lead contact

Further information and requests for resources and reagents should be directed to and will be fulfilled by the lead contact, Bradford B. Lowell.

#### Materials availability

Knock-in mouse lines will be deposited at Jackson Laboratory.

### EXPERIMENTAL MODEL AND STUDY PARTICIPANT DETAILS

#### Animals

All animal care and experimental procedures were approved in advance by the National Institutes of Health and the Beth Israel Deaconess Medical Center Institutional Animal Care and Use Committee. Mouse health was monitored daily, and mice were housed in a humidity– and temperature-controlled environment (22-24°C) with a 12 h light / 12 h dark cycle and *ad libitum* access to standard chow (LabDiet 5001) and water, unless otherwise specified. All lines were maintained on a mixed genetic background, and all experimental mice were heterozygous for the allele(s) of interest. For experiments involving across-animal comparisons, littermates of the same sex were randomly assigned to experimental or control groups. Both male and female mice were used for all studies. The following mouse lines were obtained from Jackson Laboratory: *Npy-IRES-FLPo* (#030211), *Lepr-IRES-Cre* (#032457). The following mouse lines have been described previously: *Npy-hrGFP*^72^, *Agrp-IRES-Cre*^56^, *Trh-IRES-Cre*^26^, *Pacap-IRES-Cre*^26^, *Trh-2a-Dre*^28^. *Ucn3-Cre* was obtained from MMRRC (#032078-UCD) through Catherine Dulac.

#### Generation of mouse lines

*Sim2-2a-Cre* and *Sim2-2a-FLPo* mouse lines were generated using Easi-CRISPR^73^. A *p2a-Cre* or *p2a-FLPo* cassette was inserted immediately upstream of the stop codon in the *Sim2* gene. To achieve this, fertilized eggs from FVB dams were injected with CAS9 protein (PNA Bio, CP01), sgRNA, and single-stranded DNA template (ssDNA) containing the *p2a-recombinase* cassette in addition to 100nt 5’ and 3’ homology arms. ssDNA sequences can be found in Figures S2 and S3. ssDNA was synthesized by Genscript (*Sim2-2a-Cre*) or IDT (*Sim2-2a-FLPo*). sgRNA was synthesized by Genscript, with the following sequence:

AGGGCCAGUCCCGCUCGGCA

Samples from the resultant offspring were screened for cassette insertion by PCR, using two pairs of primers targeting short amplicons straddling the 5’ and 3’ ends of the inserted cassette. Dimethyl sulfoxide (DMSO, 5%) was added to PCR reactions due to high G/C content. Positive samples were then amplified using two pairs of primers that each target a ∼1kb amplicon straddling the 5’ or 3’ end of the inserted cassette, with the two amplicons overlapping in the middle of the cassette. The resultant cDNA samples were verified for sequence integrity using Sanger sequencing (Eton Biosciences). One founder mouse was selected for each line and bred to wild-type mice of mixed background.

Primer pairs used for screening offspring were as follows:

*Sim2-2a-Cre*:

Short, left arm: TCGTGCTGCTCAACTACCAC and CATGTCCATCAGGTTCTTGC

Short, right arm: CGCTGGAGTTTCAATACCGG and CCAACCCTCTACACCAGCAT

Long, left arm: TCGTGCTGCTCAACTACCAC and AACCAGCGTTTTCGTTCTGC

Long, right arm: AGCCGAAATTGCCAGGATCA and CCAACCCTCTACACCAGCAT *Sim2-2a-FLPo*:

Short, left arm: TCGTGCTGCTCAACTACCAC and TCGAACTGGCTCATCACCTT

Short, right arm: GAGCAGCTACATCAACAGGC and CCAACCCTCTACACCAGCAT

Long, left arm: TCGTGCTGCTCAACTACCAC and ACTGAATGATCACGCCCAGG

Long, right arm: CCACATTCATCAACTGCGGC and CCAACCCTCTACACCAGCAT

### METHOD DETAILS

#### Single molecule fluorescent *in situ* hybridization (RNAscope)

Mice were terminally anaesthetized using 7% chloral hydrate (500mg kg^-1^; Sigma Aldrich) diluted in saline (0.9%), and transcardially perfused, first with 0.1 M phosphate-buffered saline (PBS), then with 4% paraformaldehyde (PFA). Brains were extracted and post-fixed overnight at 4°C in 4% PFA, and then cryopreserved in 15% and 30% sucrose for one day each at 4°C. The following day, brains were cut into 40 µm sections on a freezing microtome (Leica Biosystems). The sections were washed in PBS containing 0.5% Triton X-100, mounted on microscope slides (Fisherbrand Superfrost Plus), and dried at room temperature (RT) for one hour. Sections were then fixed for 15 minutes in 4% PFA at 4°C, washed in PBS, dehydrated in an ethanol series, and then air-dried. A barrier was drawn around the sections using an ImmEdge Hydrophobic Barrier Pen. The sections were then incubated in Protease III for 30 minutes at RT, then incubated with target probes: C3-Sim2 (Cat# 1110401-C3), C2-Trh (Cat# 436811-C2), C1-Adcyap1 (Cat# 405911), C1-Slc17a6 (Cat# 319171), C2-Bdnf (Cat# 424821), and C1-Cbln2 (Cat# 428551) for two hours at 40°C. Slides were stored overnight at 4°C in 5x SSC before being treated with Amp 1-3 at 40°C. Probes were visualized using HRP-C1/2/3 and TSA-fluorophores: Cf488a, Cf568, and Cf640r (Biotium). Sections were counterstained with DAPI, coverslipped with mounting medium (SouthernBiotech #0100-01) and imaged using a confocal microscope (Zeiss LSM 880). DAPI-labelled cells containing each of the probes were counted manually using QuPath or FIJI.

#### Stereotactic surgical procedures

Stereotactic injections were performed as previously described^5^. Mice were anesthetized by injection of xylazine (10 mg kg^-1^) and ketamine (100 mg kg^-1^) diluted in sterile saline (0.9%), and their heads fixed into a stereotaxic apparatus (Kopf). A skin incision was used to expose the skull, and a small drill hole was made above the injection site. AAVs were injected using a pulled glass micropipette (20-40 µm tip diameter) at the following coordinates (from Bregma):

PVH – AP: –0.70, ML: ±0.2, DV: –4.85

ARC – AP: –1.65, ML: ±0.3, DV: –5.90 and –5.80

vDMH – AP: –1.80, ML: ±0.3, DV: –5.20

BNST – AP: +0.40, ML: –0.55, DV: –4.15

Virus delivery was controlled by an air pressure system using picoliter air puffs through a solenoid valve (Clippard EV 24VDC) pulsed by a Grass S48 stimulator. The pipette was removed 5 minutes post-injection, and the incision was closed using VetBond (unless an optic fiber was also implanted – see below). Post-operative analgesia was provided by subcutaneous injection of sustained-release Meloxicam (4 mg kg^-1^). All subjects determined to be “misses” based on little or absent reporter expression at the target site upon histological examination were excluded from analyses.

#### Viruses

The viruses were injected at the following specified volumes (per hemisphere). The following viruses were obtained from Addgene: AAV9-CAG-Flex-rev-ChR2-tdTomato (#18917 – 35nl, unilateral), AAV5-Syn-Flex-rc[ChrimsonR-tdTomato] (#62723 – 35nl, unilateral), AAV8-EF1α-fDIO-GCaMP6s (#105714 – PVH: 35nl; ARC: 200nl per site, unilateral), AAV8-hSyn-DIO-hM3Dq-mCherry (#44361 – 35nl, bilateral), AAV1-Syn-Flex-GCaMP6s (#100845 – 200nl per site, unilateral), AAV8-hSyn-DIO-hM4Di-mCherry (#44362 – 50nl, bilateral) and AAVretro-EF1α-fDIO-Cre(HA) (#121675 – BNST: 50nl, unilateral; ARC: 75nl at –5.90 DV and 25nl at –5.80 DV). The following viruses were obtained from the Stanford Gene Vector and Virus Core (RRID:SCR_023250): AAVDJ-CMV-DIO-eGFP-2a-TeNT (GVVC-AAV-71 – 50nl, bilateral) and AAVDJ-CMV-DIO-eGFP (GVVC-AAV-12 – 50nl, bilateral). AAV8-EF1α-DIO-Syp-mCherry (35nl, unilateral) was obtained from the MGH Vector Core Facility, Charlestown, MA, USA.

**AAV8-nEF-Cre^ON^Dre^ON^-hM3Dq-mCherry** was generated by first inserting FREX from an AAV-CAG-FREX vector (kindly provided by T. Badea) into a pAAV-nEF backbone^74^ at *BamHI* and *EcoRI* sites. We designed an hM3Dq-mCherry sequence split into 3 exons with synthetic introns, and where the central exon was double-floxed in reverse orientation, following the approach of ^74^. This split-hM3Dq-mCherry sequence was inserted into the vector in reverse orientation between the double inverse rox12 and roxP sites at *AscI* and *NheI* sites. We verified that the resultant AAV drove mCherry expression in the presence of both Cre and Dre, but no expression in the presence of Cre alone, and minimal expression in the presence of Dre alone (Figure S1B).

#### Optic fiber implants

For fiber photometry experiments, following AAV injection, an optic fiber (400 µm core diameter, 0.5 NA, multimode, Thorlabs) affixed into a metal ferrule (Precision Fiber Products) was implanted 100 µm dorsal to the AAV injection site (ARC – AP: –1.65, ML: –0.3, DV: –5.75; PVH – AP: –0.70, ML: –0.2, DV: –4.75; vDMH: AP: –1.80, ML: –0.3, DV: –5.10). The skull and ferrule were scored to improve adhesion, and the ferrule was affixed to the skull using dental acrylic mixed with glue (Krazy Glue). Fiber tip location was determined upon post-mortem histological examination.

#### Chemogenetic food intake studies

Experimental mice were injected with AAV driving expression of excitatory (hM3Dq) or inhibitory (hM4Di) chemogenetic receptors in the PVH. At least 2 weeks later, animals were subjected to sham intraperitoneal (IP) injections and extensive handling every day for at least 1 week prior to food intake studies. All food intake studies were conducted starting at ZT3 (9am), when *ad libitum* fed mice have low drive to feed. Each animal was subjected to at least one trial with IP vehicle (1% DMSO in 0.9% saline) injection, and the same number of trials with IP CNO (1 mg kg^-1^ in 1% DMSO, 0.9% saline) injection. Where more than one trial was performed per condition, the mean over trials was calculated for each mouse.

For experiments involving hM3Dq, mice were moved to a new cage with food the preceding afternoon to minimize the amount of food in bedding. The next morning, 15 minutes prior to the start of the food intake study, vehicle or CNO was injected, and food was removed from the cage. Food was placed in the hopper at ZT3 and weighed every hour for 3 hours. For experiments with AAV8-nEF-Cre^ON^Dre^ON^-hM3Dq-mCherry, food was weighed at 8 hours.

For experiments involving hM4Di, mice were fasted by moving to a new cage without food at ZT12 the preceding day. Vehicle or CNO was injected at ZT2 (i.e., one hour prior to the introduction of food. Food was placed in the hopper at ZT3, and weighed every hour for 3 hours.

#### Immunofluorescence

Mice were terminally anesthetized using 7% chloral hydrate (500mg kg-1; Sigma Aldrich) diluted in saline (0.9%), and transcardially perfused, first with 0.1 M phosphate-buffered saline (PBS), then with 10% neutral-buffered formalin (NBF; Thermo Fisher Scientific). Brains were extracted and post-fixed overnight at 4°C in 10% NBF, and then cryopreserved overnight at 4°C in 20% sucrose. Brains were then cut into 40-50 µm coronal sections using a freezing microtome (Leica Biosystems) and stored at 4°C in PBS until staining. Subsequent steps were performed at room temperature. Sections were washed (4 x 3 minutes) in PBS, and then incubated for 1 hour in blocking solution (5% normal donkey serum and 0.5% Triton X-100 in PBS). Sections were then incubated overnight with the following primary antibodies diluted in blocking solution: Rat anti-mCherry (Invitrogen #M11217, 1:3000), Rabbit anti-GFP (Invitrogen #A11122, 1:3000), Rabbit anti-fluorogold (Millipore #AB153-I, 1:400) and/or Rabbit anti-c-Fos (Synaptic Systems #226-003 [since discontinued], 1:1000). The next day, sections were washed (4 x 3 mins) in PBS, then incubated for 1-2 hours with the following Alexa Fluor-conjugated Donkey secondary antibodies at 1:1000 concentration in blocking solution: anti-rat 594 (Invitrogen #21209), and anti-rabbit 488 (Invitrogen #A21206). Finally, sections were washed (4 x 3 mins) in PBS, mounted on gelatin-coated slides, air-dried, mounted using mounting medium (SouthernBiotech #0100-20) containing DAPI, and covered with a coverslip. Fluorescent images were captured using an Olympus VS120 slide-scanning microscope.

#### Thick tissue immunofluorescence

700 µm-thick coronal sections through the hypothalamus were obtained using a microtome (Leica VT1000S). Sections were cleared using CUBIC Trial Kit (Code No. 290-80801)^37^ and processed as in^75^. Briefly, sections were immersed in ScaleCUBIC-1 diluted 1:1 with MQ-H_2_O overnight in a 37°C incubator on a shaker. The tissue was then placed in ScaleCUBIC-1 for two days in a 37°C incubator on a shaker. Once the sections were sufficiently cleared, they were washed overnight at RT in PBS + 0.2% Triton X-100. Next, we incubated the sections with the primary antibody (Rat anti-mCherry, Invitrogen #M11217, 1:500) diluted in modified blocking solution (PBS with 10% Triton X-100, 5% normal horse serum and 300mM NaCl) in a 37°C incubator on a shaker for 4 days. Sections were washed overnight at RT in PBS + 0.2% Triton X-100 before being incubated in the secondary antibody (anti-rat 594, Invitrogen #21209, 1:500) and DAPI in the same blocking solution as above in a 37°C incubator on a shaker for 4 days. The sections were then washed overnight at RT in PBS + 0.2% Triton X-100 before being gently post-fixed in 1% PFA in PBS for 1 hour at RT before being washed overnight at RT in PBS + 0.2% Triton X-100. Sections were then immersed in ScaleCUBIC-2 diluted 1:1 with MQ-H_2_O overnight in a 37°C incubator on a shaker, before being immersed in ScaleCUBIC-2 overnight in a 37°C incubator on a shaker. Sections were imaged in a 2:8 mix of mounting solution 1 and 2 (CUBIC Trial Kit, Code No. 290-80801) to achieve a refractive index of ∼1.49. Sections were imaged using a Leica SP5 confocal microscope at 10x magnification.

#### Fluorogold labelling

We used peripheral injection of fluorogold to label neuroendocrine neurons that project outside the blood-brain barrier (BBB), since fluorogold cannot cross the BBB, but is taken up by axon terminals outside the BBB and transported retrogradely to label cell bodies^6^. *Sim2-2a-Cre*::*tdTomato* mice were injected intraperitoneally with fluorogold (50µl of a 5% solution in 0.9% NaCl, Fluorochrome), and mice were perfused as described above 5 days later. Sections from each mouse were matched at 4 anterior-posterior levels of PVH, visualized using immunofluorescence, and imaged using a confocal microscope (Zeiss LSM 880). Cell bodies labelled with *tdTomato* and/or fluorogold were counted manually using QuPath.

#### Collateral mapping

To map collaterals of ARC– and BNST-projecting PVH^Sim2^ neurons, *Sim2-2a-FLPo* mice were injected with AAVretro-EF1α-fDIO-Cre(HA) in the ARC or BNST, respectively, and AAV8-EF1α-DIO-Syp-mCherry in the PVH. Animals were perfused 3 weeks after injection. The presence of collaterals in BNST/ARC was determined based on immunofluorescent detection of mCherry expression.

#### Channelrhodopsin-assisted circuit mapping

*Sim2-2a-Cre::Npy-hrGFP* or *Ucn3-Cre::Npy-hrGFP* mice were injected with AAV9-CAG-Flex-rev-ChR2-tdTomato in the PVH. To obtain *ex vivo* sections containing the ARC, mice were deeply anaesthetized using isoflurane and decapitated, and the brain was removed and immediately submerged in ice-cold cutting solution (in mM: 92 choline chloride, 10 HEPES, 2.5 KCl, 1.25 Na_2_HPO_4_, 30 NaHCO_3_, 25 glucose, 10 MgSO_4_, 0.5 CaCl_2_, 2 thiourea, 5 sodium ascorbate, 3 sodium pyruvate) saturated with carbogen (95% O_2_, 5% CO_2_) and with measured osmolarity 310-320 mOsm l^-1^ and pH 7.4. Coronal sections (275-300 µm thick) were then cut using a vibratome (Campden 7000smz-2) and incubated in carbogen-saturated cutting solution for 10 minutes at 34°C. Sections were then transferred to oxygenated artificial cerebrospinal fluid (aCSF – in mM: 126 NaCl, 21.4 NaHCO_3_, 2.5 KCl, 1.2 NaH_2_PO_4_, 1.2 MgCl_2_, 2.4 CaCl_2_, 10 glucose) at 34°C for 15 minutes. Sections were then kept in oxygenated aCSF at room temperature until use. For recordings, a single section was placed into the recording chamber, where it was continually perfused with oxygenated aCSF at 3-4 ml min^-1^. Neurons were visualized using an upright microscope (SliceScope Pro 1000, Scientifica) equipped with infrared differential interference contrast and fluorescence optics. Patch pipettes with open-tip resistances of 3-5 MΩ were backfilled with internal solution (in mM: 135 CsMeSO_3_, 10 HEPES, 1 EGTA, 3.3 QX-314 (Cl^-^ salt), 4 Mg_2_-ATP, 0.3 Na_2_GTP, and 8 Na_2_-phosphocreatine) with measured osmolarity 290 mOsm l^-1^ and pH 7.35. Optogenetically-evoked excitatory post-synaptic currents (oEPSCs) were isolated in whole-cell voltage clamp at V_h_=-70 mV. To photostimulate ChR2-expressing fibers in ARC, blue light from an LED (470 nm, Cool LED pE-100) was focused onto the back aperture of the microscope objective (40x) to produce wide-field illumination around the recorded neuron at a power of 10-15 mW mm^-2^ (as measured using an optical power meter, PM100D, Thorlabs). The blue LED was controlled by a programmable pulse stimulator (Master 8, A.M.P.I.) and pClamp 10.5 software (Axon Instruments). The stimulation protocol consisted of four blue light pulses (470 nm wavelength, 5 msec) administered 1 s apart during the first 4 s of an 8-s sweep, repeated for a total of 20 sweeps. EPSCs with a latency of ≤6 ms following light delivery were considered as likely monosynaptic oEPSCs. To confirm the monosynaptic nature of PVH^Sim2^→ARC^AgRP^ connections, for a subset of recorded neurons, oEPSCs were recorded in the presence of tetrodotoxin (TTX, 1 µM) alone and then in combination with 4-aminopyridine (4-AP, 500 µM). Signals were amplified through a Multiclamp 700B amplifier, and data were filtered at 2 kHz and digitized at 20 Hz. Access resistance (<30 Ω) was continuously monitored by a voltage step, and recordings were used for analysis if resistance changed by <15%. All recordings were analyzed using Clampfit 10.5.

#### Fos studies

*Sim2-2a-FLPo*::*tdTomato* mice were single-housed and habituated to handling for at least one week prior to the start of the experiment. Then, cages were changed at ZT10, and no food was introduced into the new cage of the fasting group. The next day at approximately ZT6, mice were anesthetized and perfused as described above. Brains were cut into 40 µm sections, and for each mouse, 4 sections were selected at matched intervals spanning the anterior-posterior extent of the PVH for manual quantification. Counts of tdTomato– and Fos-expressing neurons were summed across the 4 sections for each mouse.

#### Hormone measurements

Wild-type mice were habituated to extensive handling for at least 10 days prior to sacrifice. Then, these mice were placed into a fresh cage with food around ZT6. At ZT12, mice in the “ZT12 *ad lib*” condition were sacrificed; food was removed from mice in the “ZT13 fast” condition, and for mice in the “ZT13 *ad lib*” condition, the cages were opened briefly but food was not removed. Then at ZT13, the remaining mice were sacrificed. All mice were sacrificed by rapid decapitation, and trunk blood was collected into K2EDTA tubes on ice.

Collected trunk blood was centrifuged, and plasma was collected into aliquots and stored at –80°C. For ghrelin measurements, the K2EDTA collection tubes contained 4 µL of 100 mM p-hydroxymercuribenzoic acid in potassium phosphate buffer (for a final concentration of 1 mM); and plasma was mixed with 1 N hydrochloric acid to reach a final concentration of 0.1 N. Plasma aliquots were then run in duplicate for each test.

Enzyme-linked immunosorbent assay (ELISA) kits were used to measure ghrelin (Millipore, EZRGRA), insulin (Crystal Chem, 90080), and leptin (R&D Systems, MOB00B) in 96-well plates, according to the manufacturer’s instructions. Absorbance was measured using a plate reader.

#### Fiber photometry

Fiber photometry acquisition was controlled using a pyPhotometry microcontroller^76^, and excitation LEDs (405 nm and 465 nm) and optical paths were contained within a fluorescence minicube (Doric Lenses ilFMC4). A low-autofluorescence optic fiber cable (1m length, 400 µm diameter, NA 0.48) transmitted excitation and emission light between the sample and minicube, and was affixed to the metal ferrule on the mouse’s skull through a ceramic mating sleeve (Thorlabs) with a small dab of UV-curing glue to prevent movement relative to the sample. Emitted light (500 – 550 nm) was measured using a photodetector embedded in the minicube, and digitized by the pyPhotometry board. Excitation light power was adjusted such that emission light fell roughly in the middle of the range of the photodetector. The sample was illuminated at 10 Hz using time-division multiplexing. All recordings were performed in the home cage, and mice were allowed to habituate to patch cord attachment for at least 10 minutes prior to the start of the recording.

Prior to short-term recordings of neural responses to food and water, mice were allowed *ad libitum* access to food, or denied access overnight to either food or water, as indicated. The same mice were subjected to different conditions and stimuli to allow for within-mouse comparisons. Immediately prior to recording, both food and water were removed from the cage. After 3 minutes of baseline recording, a pellet of chow (LabDiet 5001) or bowl of water was placed into the mouse’s cage, and the recording continued for a further 24 minutes. For a subset of these recordings, video recordings of the mouse were simultaneously acquired at 10 Hz using a camera (Allied Vision Mako U-130B), with acquisition controlled by Bonsai^77^. Video and photometry data were synchronized using a TTL pulse sent by the camera to the pyPhotometry board upon each acquired frame.

For long-term recordings, mice expressing GCaMP6s in PVH^Sim2^ or ARC^AgRP^ neurons were first habituated to the recording setup by being tethered to the patch cord for at least 30 minutes per day over 2-3 days. Then, during long-term recordings, mice had access to food and water in separate bowls in their home cage, and recordings began before ZT6. For experiments involving fasting or water deprivation, food or water was manually removed from the cage at the specified time, and returned the following morning at ZT2. The recording was stopped after ZT5. Following experiments involving fasting or water deprivation, mice were allowed at least 5 days for recovery before being used for another experiment. In all studies, the same animals were subjected to conditions being compared, to allow within-animal comparisons.

For all experiments involving recording from ARC^AgRP^ neurons, each mouse was first screened for signal quality by measuring the neural response to a food pellet following overnight fasting. As previously demonstrated^7–9^, ARC^AgRP^ activity drops substantially in response to food, so mice that showed no drop in signal over 3 trials on different days were excluded from further experiments.

To record from PVH^Sim2^ neurons, *Sim2-2a-FLPo* mice expressing AAV8-EF1α-fDIO-GCaMP6s in the PVH were used. To record from ARC^AgRP^ neurons, *Agrp-IRES-Cre* mice expressing AAV1-hSyn-DIO-GCaMP6s in ARC were used. To record from ARC^AgRP^ neurons while inhibiting PVH^Sim2^ or PVH^Trh^ neurons, *Npy-IRES-FLPo*::*Sim2-2a-Cre* or *Trh-IRES-Cre* mice, respectively, expressed AAV8-EF1α-fDIO-GCaMP6s in the ARC and AAV8-hSyn-DIO-hM4Di-mCherry in the PVH. In the latter experiments, clozapine N-oxide (CNO, 1 mg kg^-1^) or vehicle (1% DMSO in 0.9% saline) was injected intraperitoneally at ZT11.5.

For experiments involving concurrent photometry recording and optogenetic stimulation, mice used were either *Sim2-2a-Cre*::*Npy-IRES-FLPo* or *AgRP-IRES-Cre*::*Sim2-2a-FLPo*, expressing AAV5-Syn-Flex-rc[ChrimsonR-tdTomato] in the PVH/ARC and AAV8-EF1α-fDIO-GCaMP6s in the ARC/PVH, respectively. 630 nm excitation light from an LED light source (Plexon) was passed through a fluorescence minicube (Doric Lenses) together with the photometry excitation light as described above, and delivered to the brain through the same optic fiber. For these experiments, photometry data was acquired using time-division multiplexed illumination at 20 Hz. Optogenetic pulses were delivered at 20 Hz, and were interleaved between the 465 nm and 405 nm photometry excitation pulses to prevent bleedthrough using custom code running on an Arduino board synchronized to the pyPhotometry system.

#### Operant task

The operant task was administered in the home cage using FED3 devices^54^ (Open Ephys) running custom Arduino code. At all stages of conditioning, mouse body weight was monitored daily, and where necessary, each step was extended until mice maintained at least their starting body weight. Mice were first habituated to 20 mg chow pellets (LabDiet 5TUM) in a bowl overnight. The next day, bedding was changed and the FED3 was introduced in free-feeding mode (i.e. mice could access pellets *ad libitum*). After one day of free-feeding, the FED3 was switched into fixed-ratio 1 (FR1; i.e. 1 nose-poke leads to pellet delivery). In FR1, both nose ports were active, and no conditioned stimulus was delivered besides the pellet. Mice spent at least 3 nights in FR1 mode, and maintained a stable body weight for at least 2 days, before photometry experiments began. For fasting, pellet delivery was paused between ZT12 and ZT2.

#### Fiber photometry data analysis

Fiber photometry data were analyzed using custom Python scripts. First, for short experiments (< 1 hour), data from both channels was filtered using a Gaussian filter (window = 7 samples, sigma = 1) to remove noise. Then, for all experiments, correction for motion artifacts was achieved by using linear regression to predict the calcium-dependent (465 nm-evoked) fluorescence based on calcium-independent (405 nm-evoked) fluorescence at each time point, subtracting this predicted signal from the measured 465 nm signal, and adding the mean measured 465 nm signal across the whole recording. Next, this corrected signal was aligned to the time of pellet drop. ΔF/F_0_ was calculated as (F-F_0_)/ F_0_ at each time point, where F_0_ is the mean signal from 3 minutes to 10 seconds prior to pellet drop. Finally, zΔF/F_0_ was calculated by taking the z-score of this signal, using the formula (F-M)/S, where F is the ΔF/F_0_ at a given timepoint; and M and S are the mean and standard deviation, respectively, of the ΔF/F_0_ signal from 3 minutes to 10 seconds prior to pellet drop.

For comparing activity before and after pellet delivery (Figure 2F), we took the mean zΔF/F_0_ in the 3 minutes before (pre) and after (post) pellet drop. For comparing the response between conditions (Figures 2H and 2J), we took the mean ΔF/F_0_ in the 5 minutes following pellet drop. For comparing different time windows relative to food contact/consumption, we took the mean ΔF/F_0_ for each trial in the following windows: pre-drop=30-20s before pellet delivery (to avoid contamination by cues before drop, such as hand over cage); pre-contact=drop to 1s before contact; pre-bite=contact to 1s before bite; bite=10s following first bite.

We calculated time constants (τ) of the food response as follows: for each mouse, within a window of 30 seconds before to 5 minutes after food presentation, we first determined the minimum signal, and then identified τ as the earliest time point at which 63.8% of this minimum had occurred, relative to the pre-stimulus baseline.

For long-term recordings (≥23 hours), motion correction was first performed as described above. Then, ΔF/F_0_ was calculated as (F-F_0_)/ F_0_ at each time point, where F_0_ is the mean signal from ZT6 to ZT9. Given our interest in slow-timescale activity changes, we then filtered out fast-timescale fluctuations by taking the rolling mean of this ΔF/F_0_ signal over a 30-minute window preceding each data point. Next, zΔF/F_0_ was calculated by taking the z-score of this filtered signal, using the formula (F-M)/S, where F is the signal at a given timepoint; and M and S are the mean and standard deviation, respectively, of the signal from ZT6 to ZT9. For recordings where fasting was initiated at ZT6, the window (ZT3-ZT6) was instead used for calculating F_0_, M, and S. Finally, for mice with more than one trial for a given condition, we took the mean over trials for each mouse, so that subsequent analyses were performed on a per-mouse basis.

For quantifying the response to food removal, we took the mean zΔF/F_0_ signal in 1-hour bins from ZT11 to ZT16 for each mouse. We calculated time constants (τ) of the response to food removal as follows: for each mouse, within a window from ZT11 to ZT18, we first determined the maximum signal, and then identified τ as the earliest time point at which 63.8% of this maximum had occurred, relative to the mean signal from ZT11 to ZT12.

For experiments with photometry recordings during the operant task, we first aligned photometry and FED3 data. Then, for each trial, we calculated the cumulative numbers of pellets retrieved and pokes (combining both nose ports). Poke rate was calculated as the mean number of pokes per minute over a sliding window of 30 minutes. To calculate activity aligned to pokes, we used only pokes during the fast period (ZT12-ZT2) that had no poke in the preceding 45 seconds. We first filtered the GCaMP6s signal (after correcting for motion artifacts – see above) using a Gaussian filter as described above for short experiments. Then, for each poke, we took this filtered signal from 45 seconds before to 100 seconds after each poke, and calculated zΔF/F_0_ as described above for short experiments, using the window of 45 s to 1 s before the poke to calculate F_0_, M, and S. Finally, we took the mean (± SEM) zΔF/F_0_ signal over all pokes from all mice. To split pokes by percentile (Figure S13), for each trial, we again used only pokes with no poke in the preceding 45 seconds. From these pokes, we calculated the cumulative percentage of pokes during the fast period at each poke time. We then binned pokes into deciles, and took the mean poke-aligned trace in each decile. For ease of visualization, we present four equally-spaced deciles.

#### Long-term body mass studies

*Sim2-2a-Cre* mice (8-13 weeks old) were injected with AAVDJ-CMV-DIO-eGFP-2a-TeNT (“TeNT”) or AAVDJ-CMV-DIO-eGFP (“GFP”) bilaterally in the PVH. Starting body weight was measured immediately prior to surgery, and following surgery mice were individually housed, and both food and body mass were weighed weekly for 12 weeks. At 13 weeks post-injection, body composition was measured using an EchoMRI 3-in-1 Analyzer without anesthesia. A single female was removed from the GFP control group for food intake studies due to evidence of food shredding.

#### Indirect calorimetry

Sim2-TeNT and Sim2-GFP mice were injected and housed as described above. At 4 weeks post-injection, mice were weighed, and their body composition analyzed as described above. They were then introduced to individual indirect calorimetry cages (Sable Systems Promethion) in the BIDMC Energy Balance Core. Mice were housed in these cages overnight (>18 h) before recording the 48 h experiment. During the experiment, the Promethion system measured physical activity (beam breaks), oxygen consumption, carbon dioxide production, body mass, and food/water consumption every 3 minutes. From these measurements, energy expenditure, respiratory exchange ratio, and total distance travelled were calculated. For non-cumulative time series (Figures S14B and S14D), the mean value was calculated over a sliding window of 30 minutes for each mouse. Analyses and statistical tests were performed using a custom Python script, except for comparing energy expenditure. Since energy expenditure can be affected by body mass, we employed a generalized linear model to account for possible effects of both treatment and body mass on energy expenditure, using the average energy expenditure over the 48 h experiment as the dependent variable^78^. This statistical model was implemented in CalR2 (https://bankslab.shinyapps.io/prod_ver/).

### QUANTIFICATION AND STATISTICAL ANALYSIS

Statistical analyses were performed using Python (for photometry and indirect calorimetry data), CalR2 (for energy expenditure measurements), or GraphPad Prism 6 or 10 (all other data), as described above. Sample sizes (N) and statistical tests are reported in figure legends. Briefly: when comparing two groups at a single time point, we used two-tailed t-tests (paired for within-subject comparisons). When comparing more than two groups, we used one-way ANOVA. When comparing groups at multiple time points, we used two-way repeated measures ANOVA. If a significant interaction (treatment*time, p<0.05) was identified, we then performed post-hoc tests at each time point, or between adjacent time points, with Holm-Sidak correction for multiple comparisons. To compare energy expenditure between groups while accounting for body mass differences, we used a generalized linear model, as described above. For all tests, α=0.05 was used as the threshold for significance.

For histological quantifications, we used QuPath or FIJI software to manually count neurons labelled by each of the specified probes/markers within the PVH at the specified anterior-posterior level(s).

---

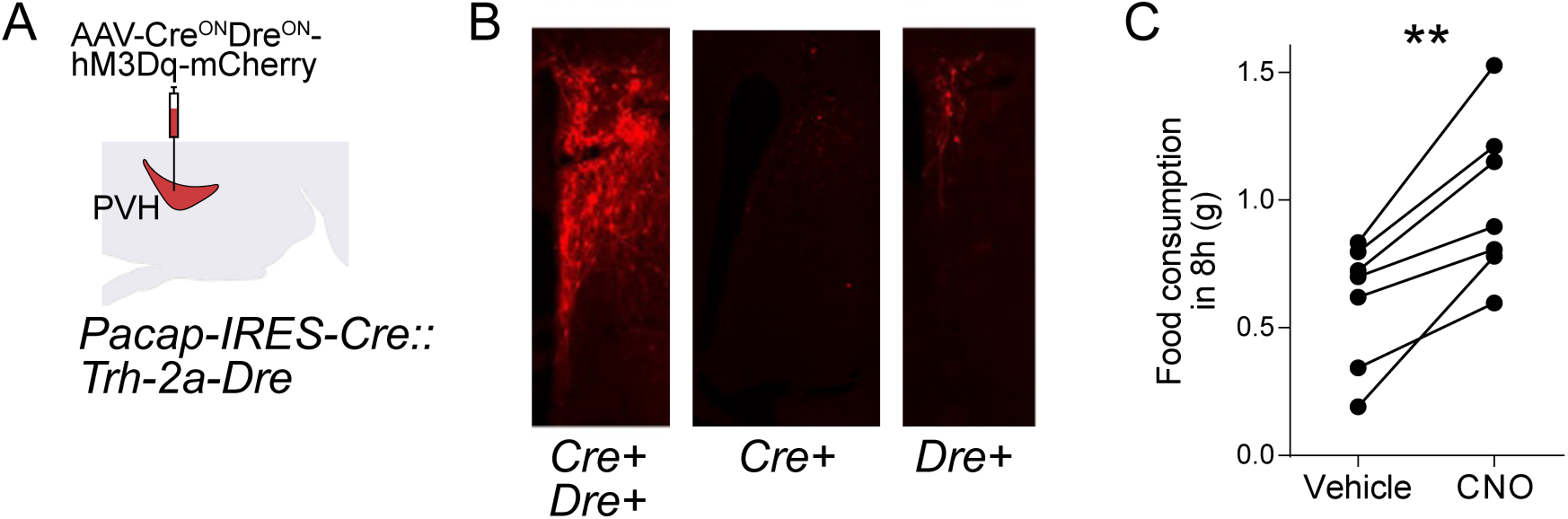
Figure S1.

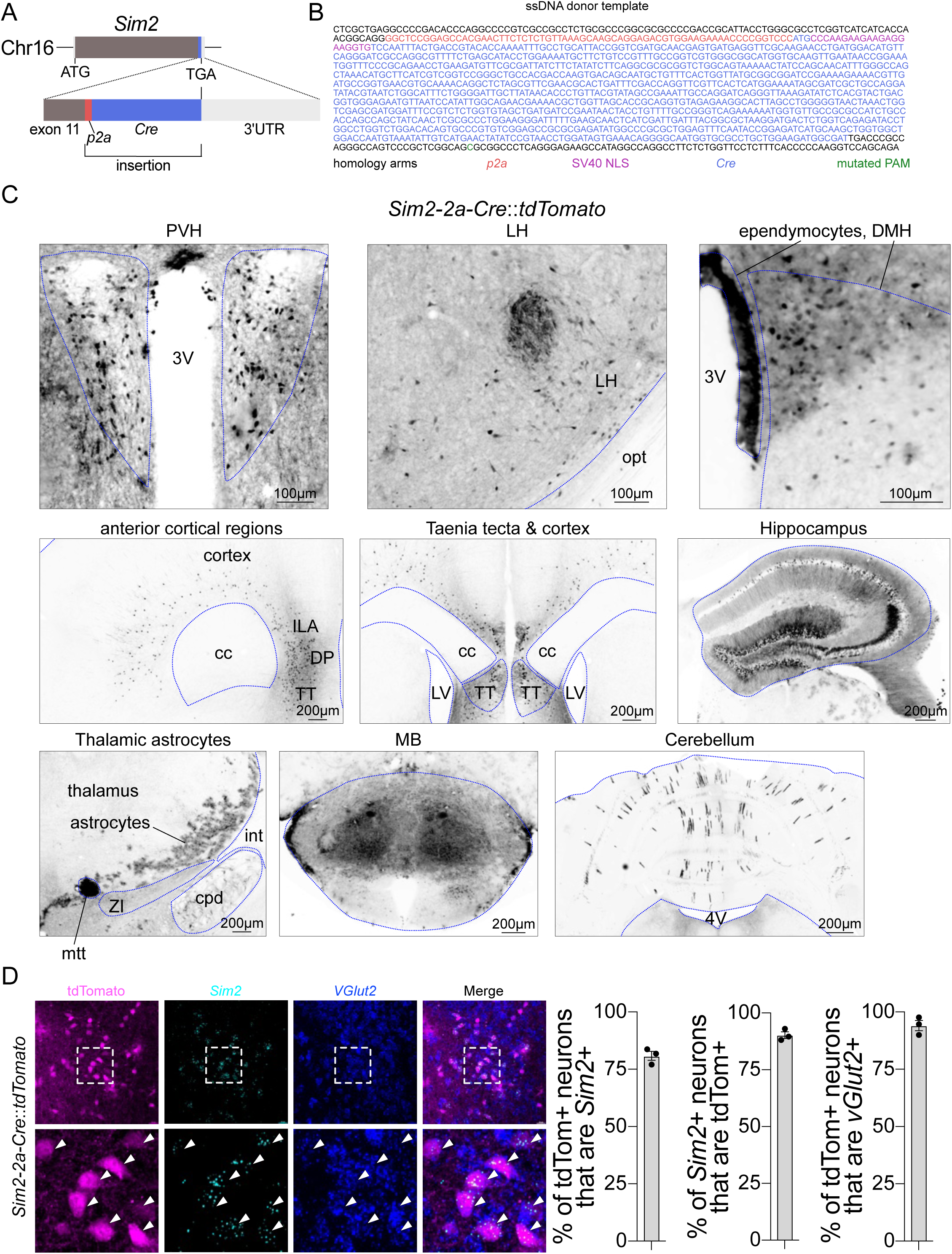
Figure S2.

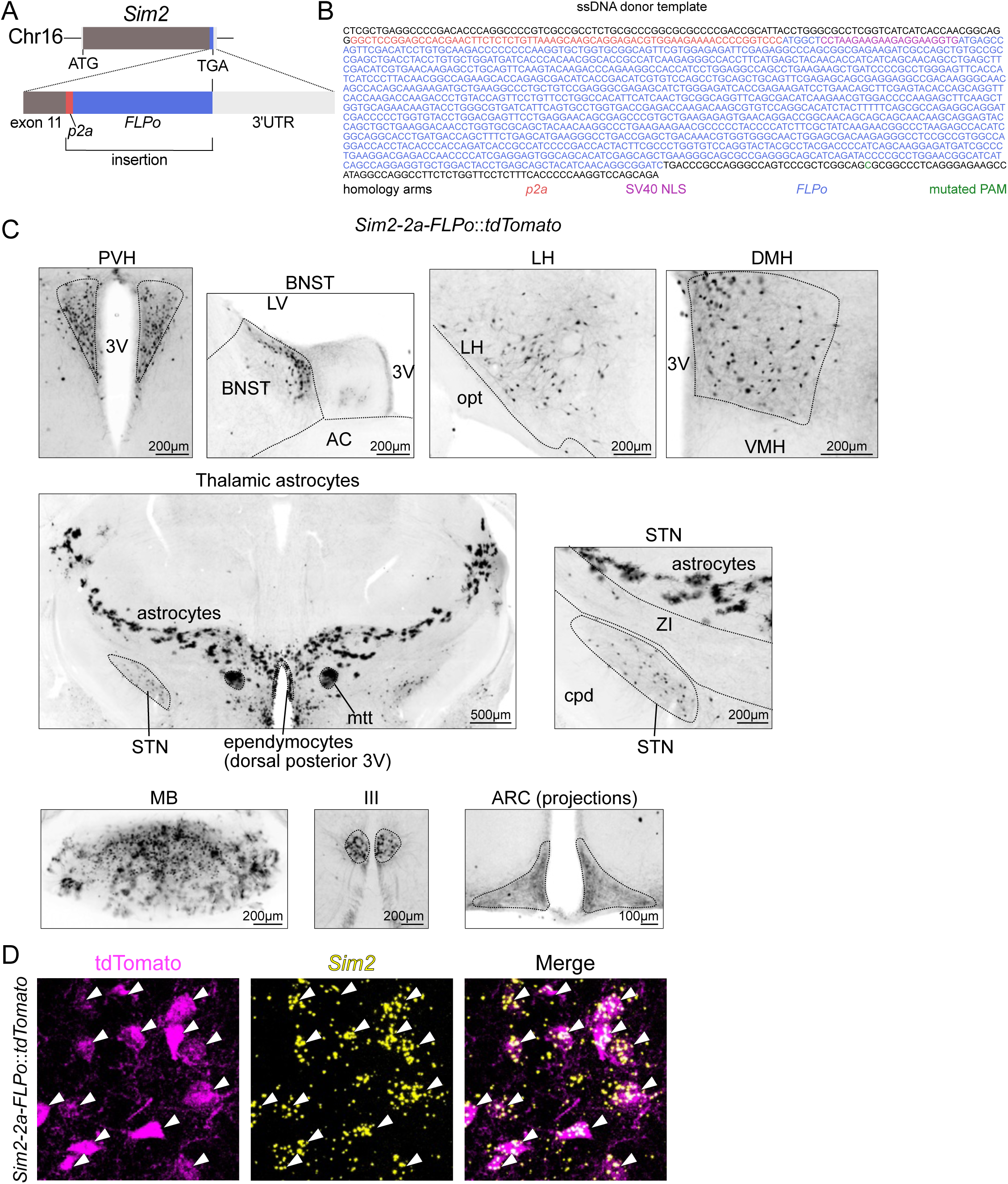
Figure S3.

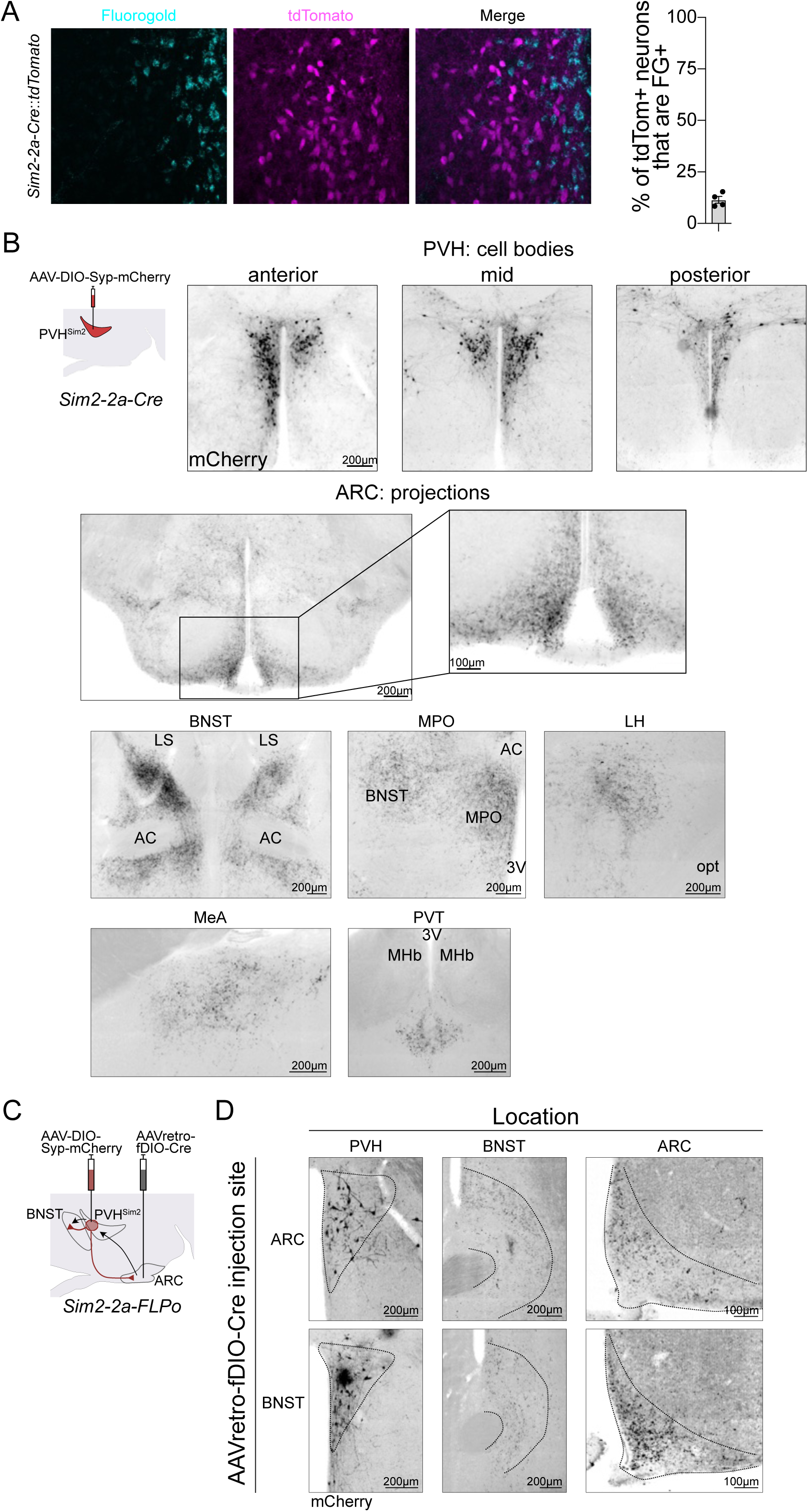
Figure S4.

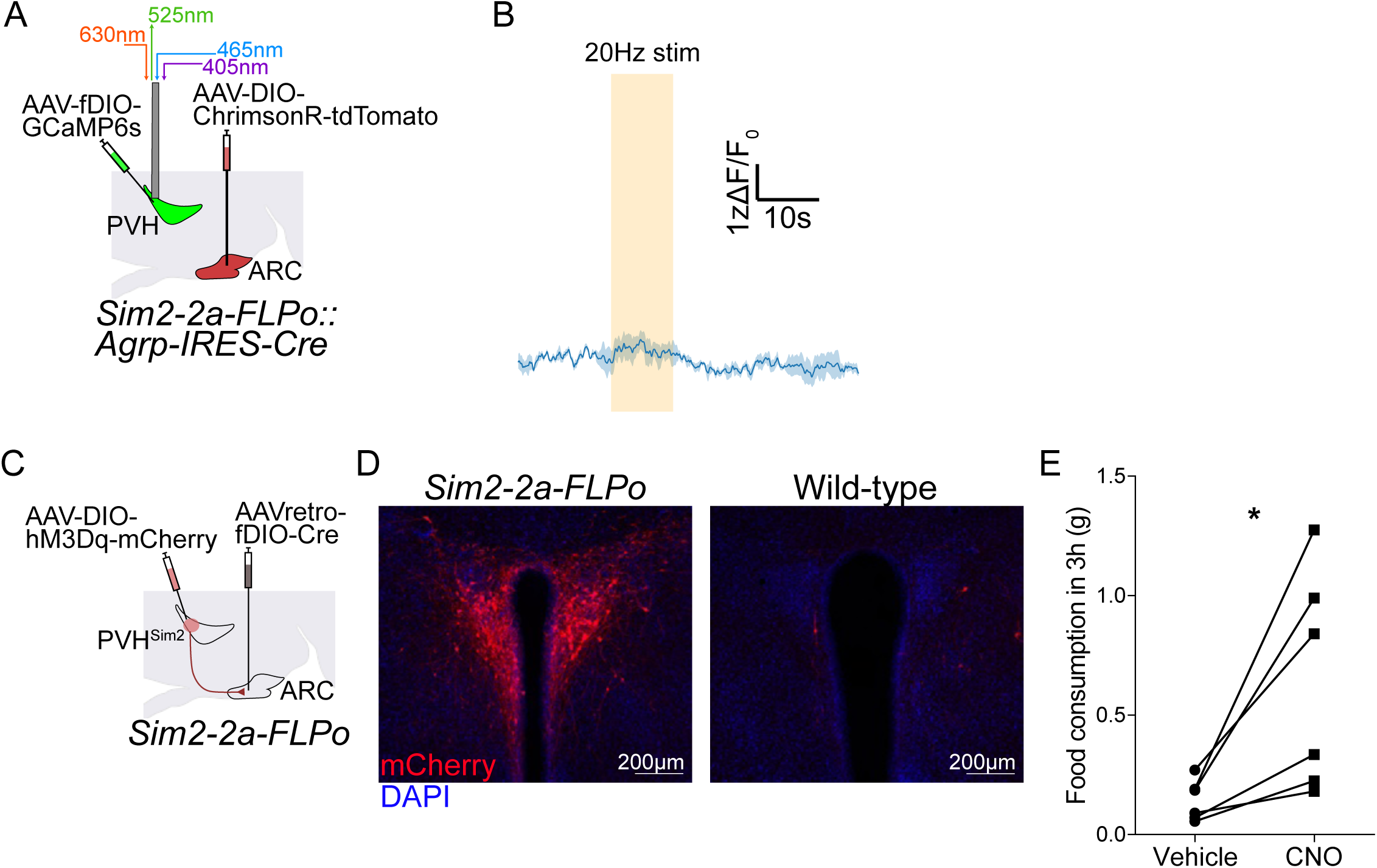
Figure S5.

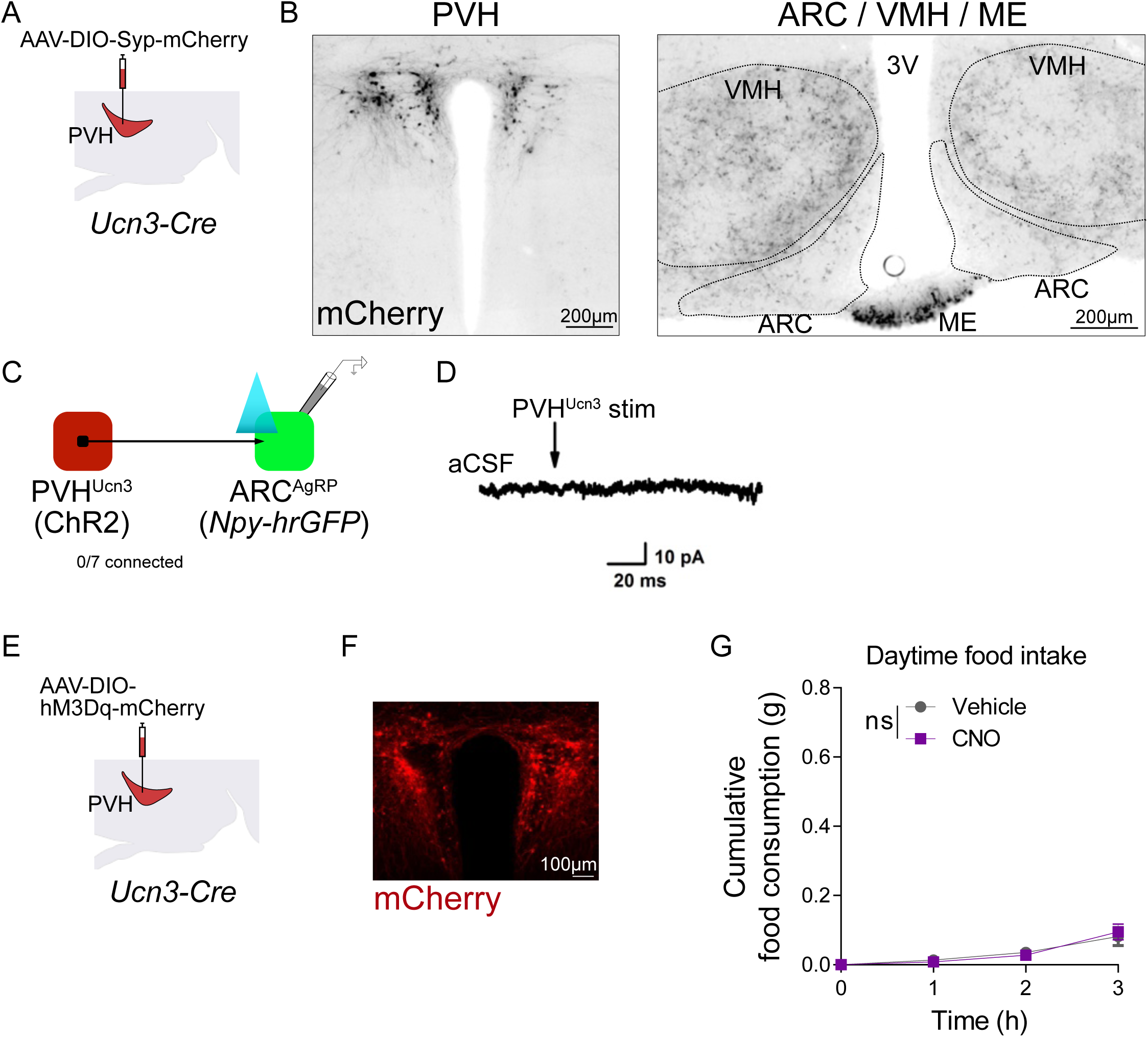
Figure S6.

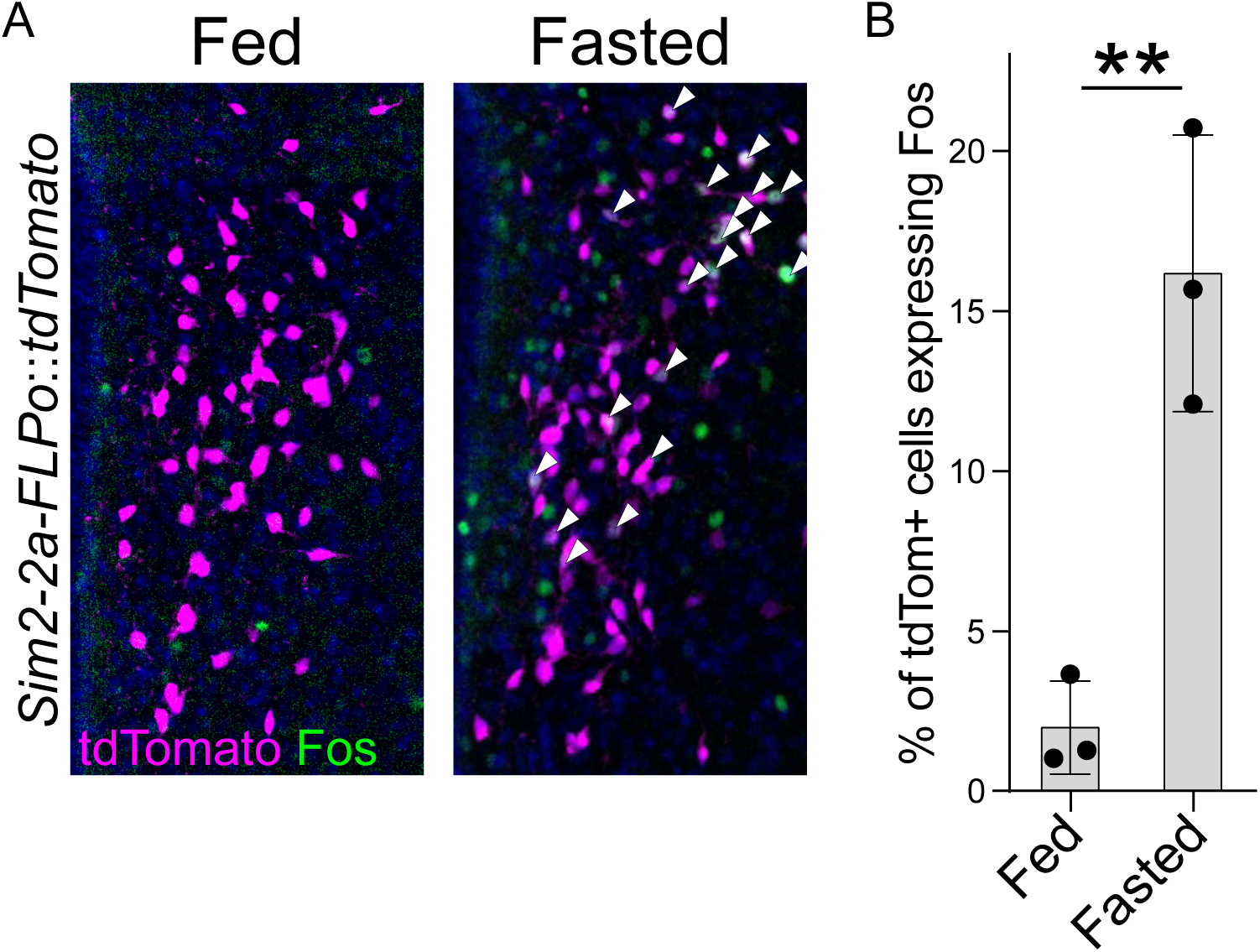
Figure S7.

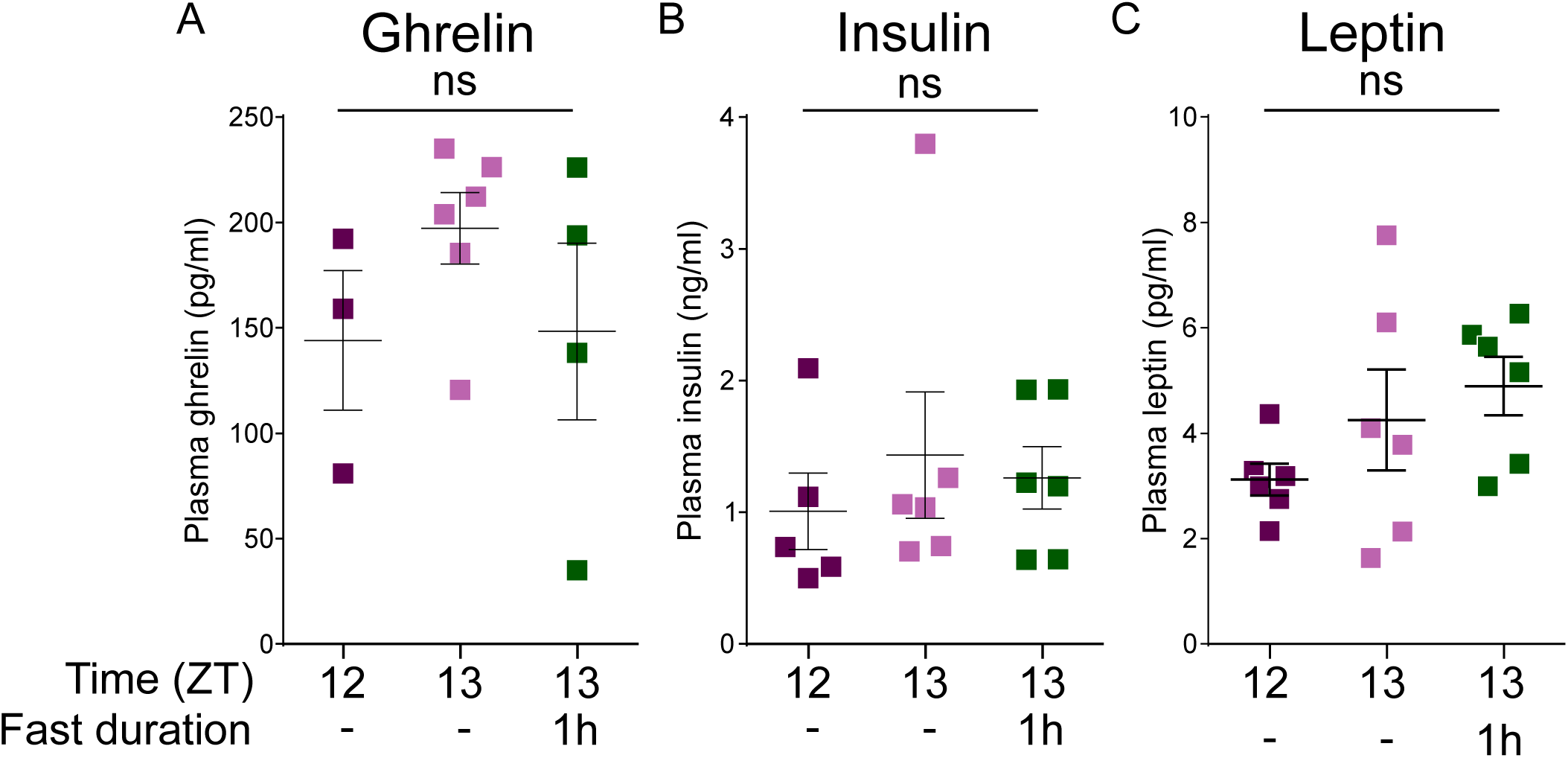
Figure S8.

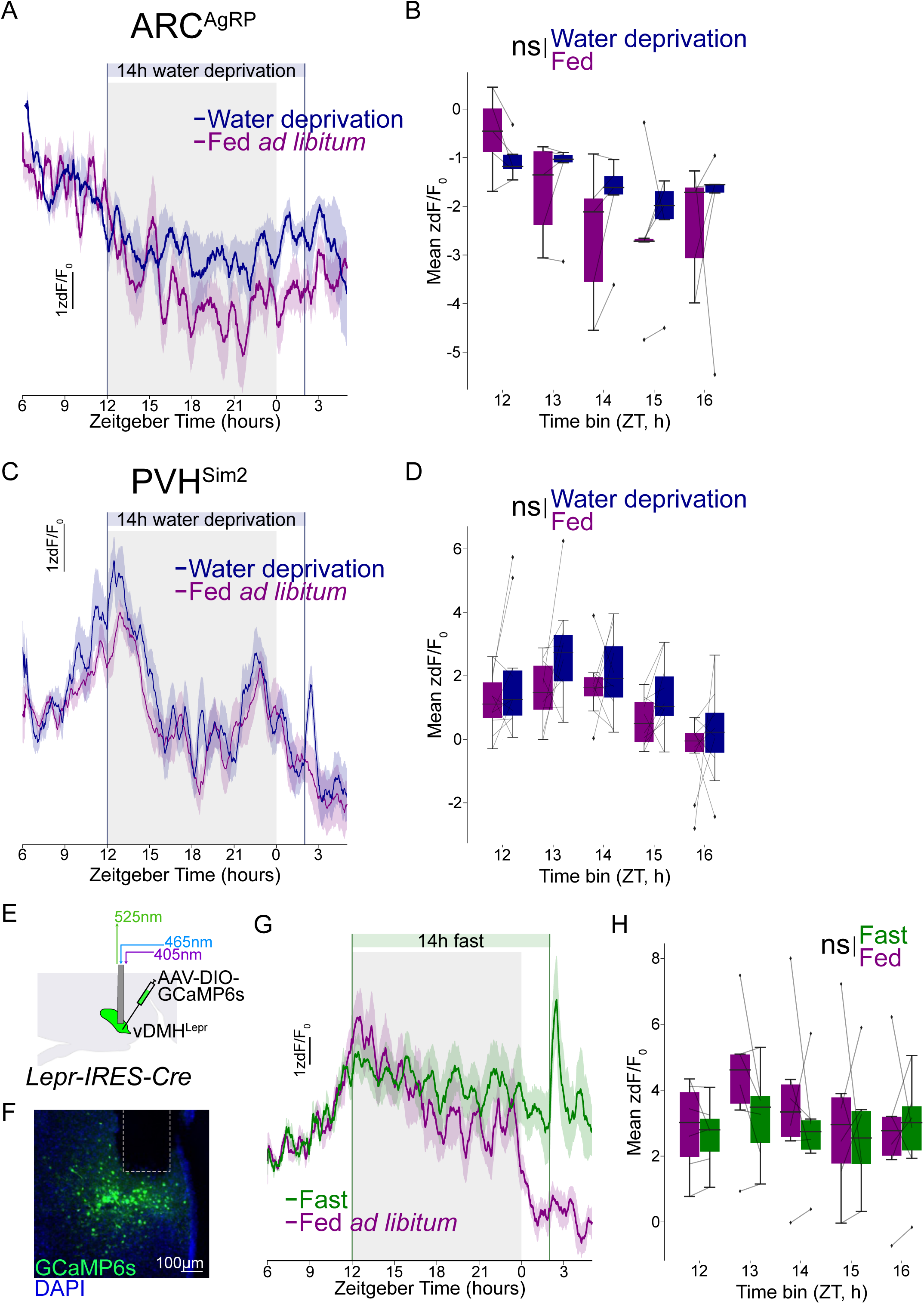
Figure S9.

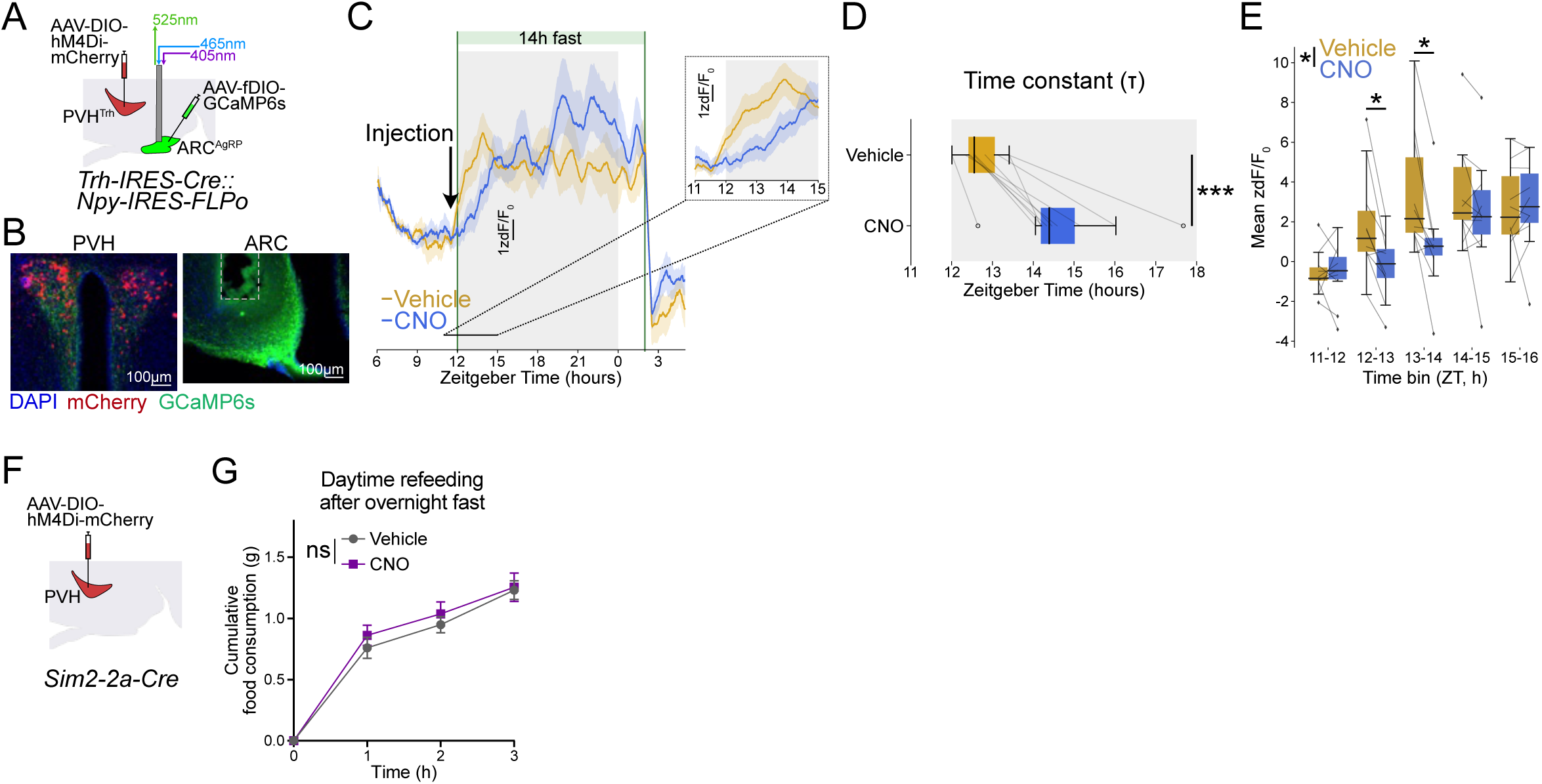
Figure S10.

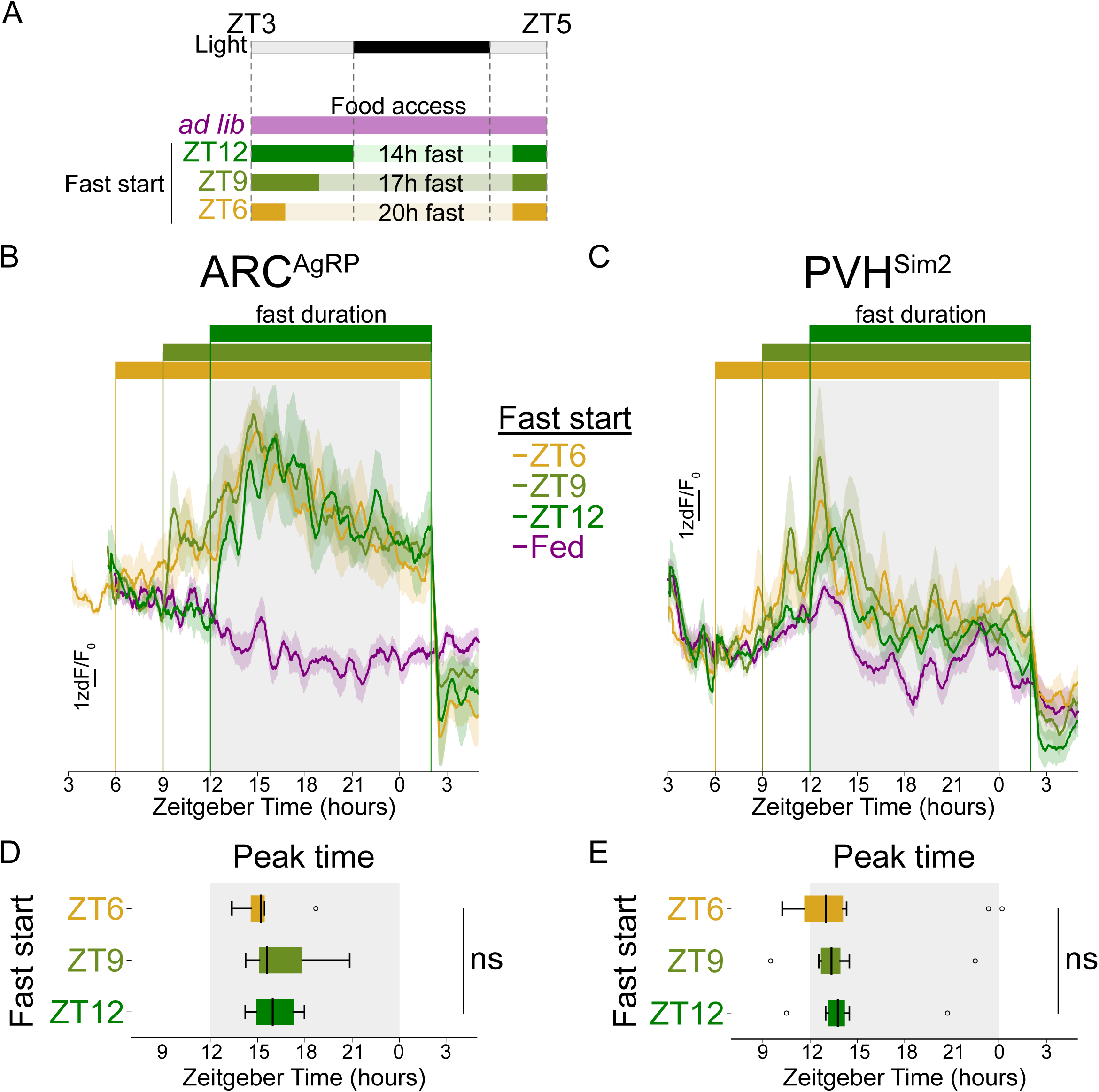
Figure S11.

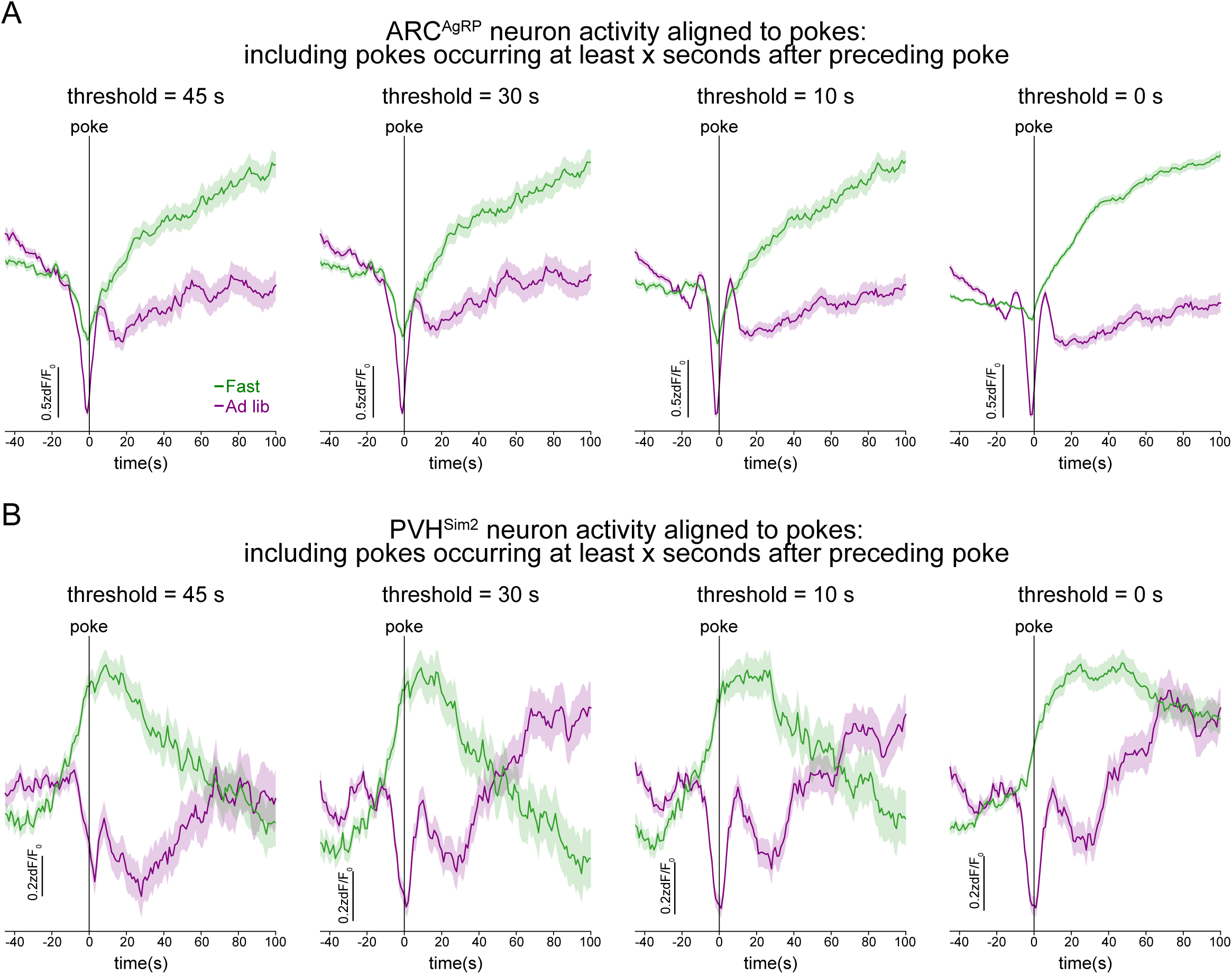
Figure S12.

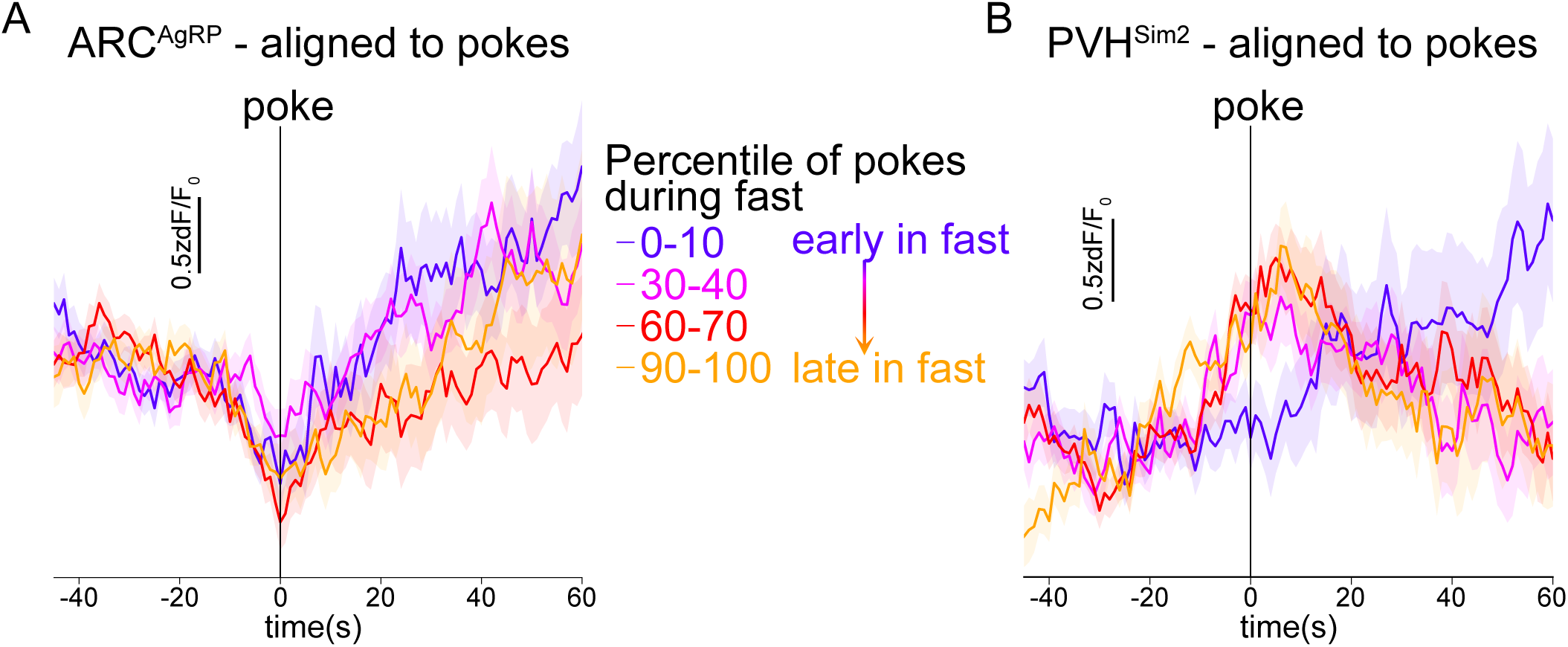
Figure S13.

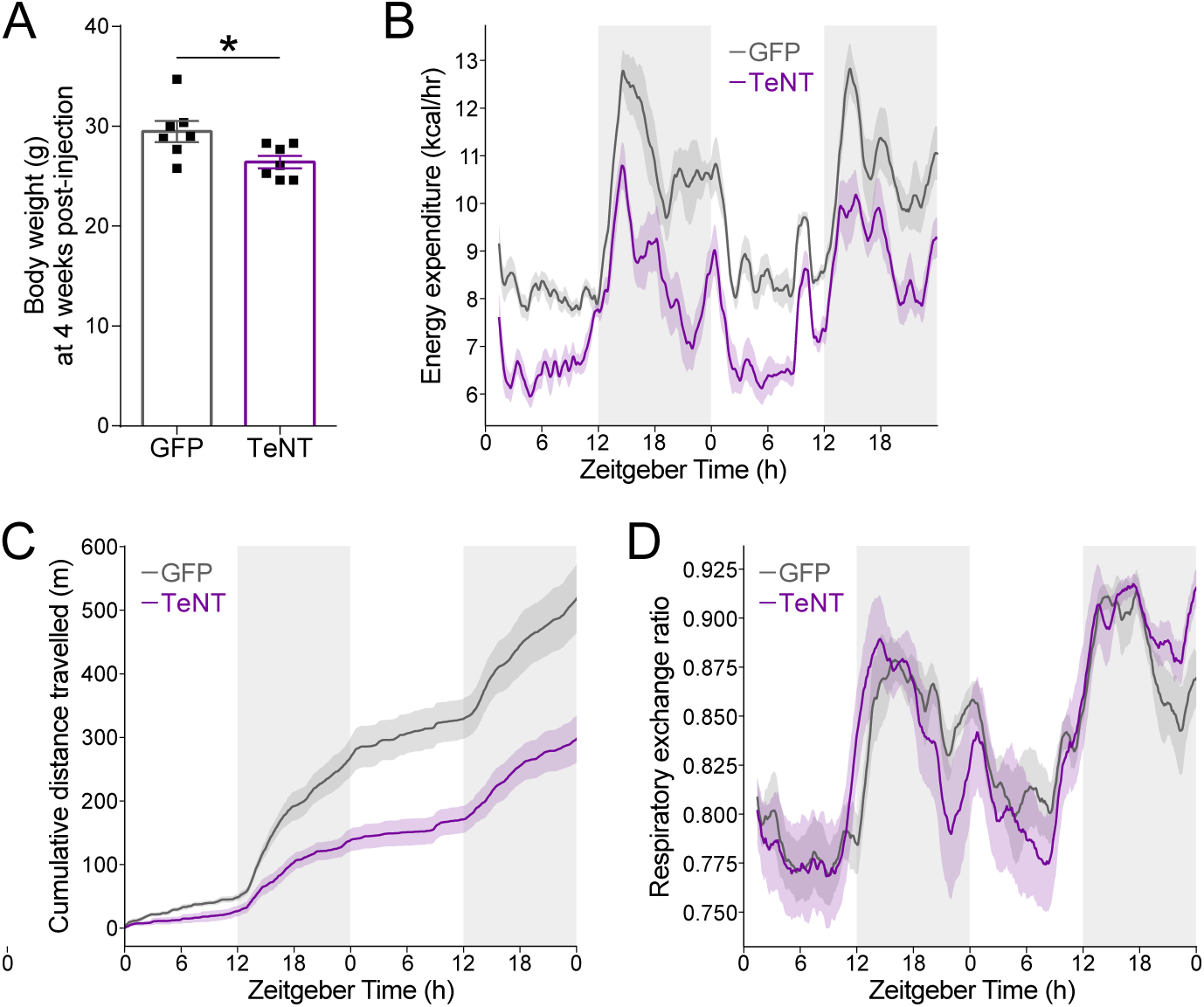
Figure S14.

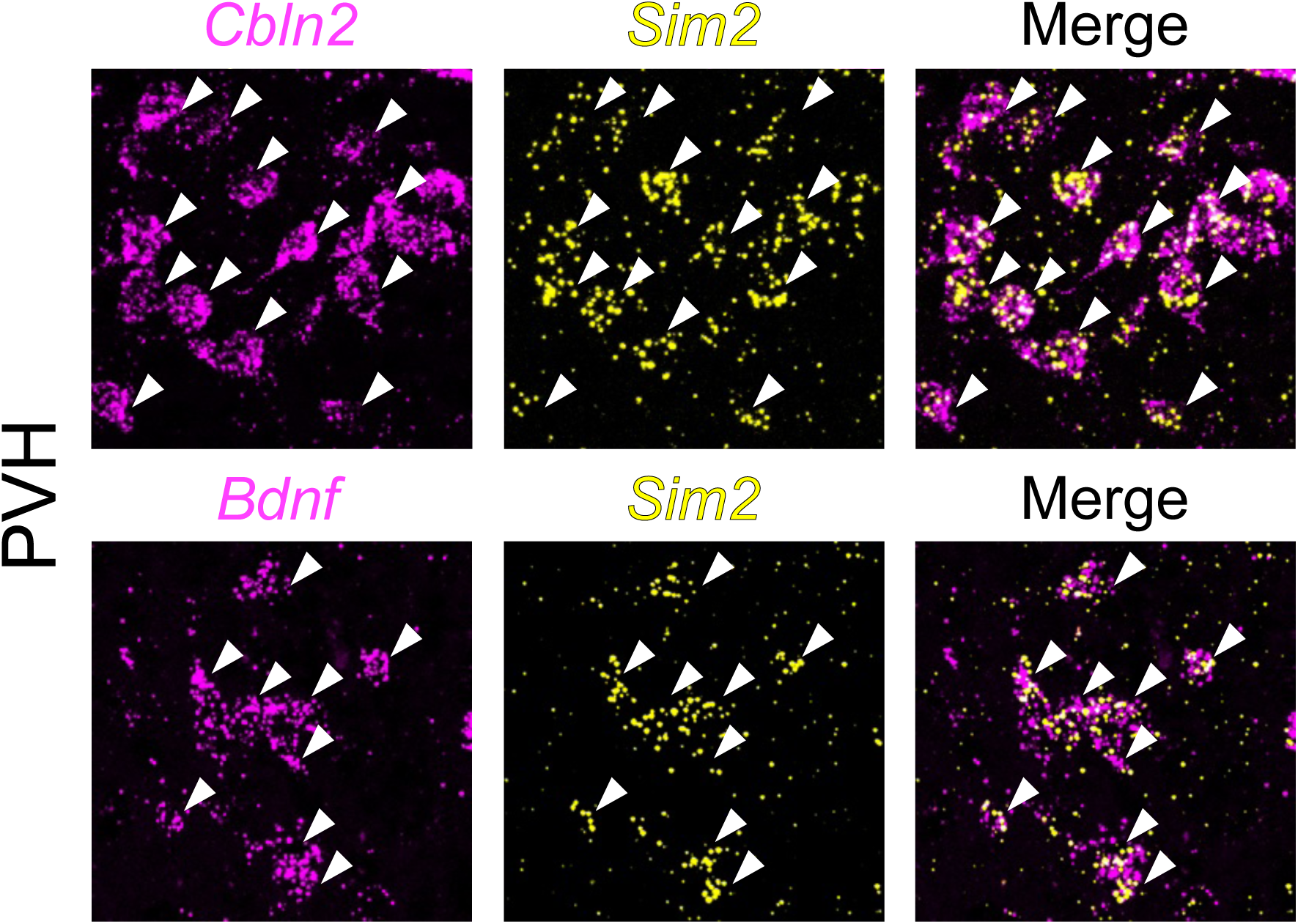
Figure S15.

