## Supplemental Figure Legends for "A hypothalamic circuit for anticipating future changes in energy balance"

**Figure S1. Activation of Trh/Adcyap1-coexpressing PVH neurons increases food intake, Related to Figure 1.**
A) Schematic of AAV-Cre$^{ON}$Dre$^{ON}$-hM3Dq-mCherry injection into the PVH of *Pacap-IRES-Cre*::*Trh-2a-Dre* mice. B) Expression of mCherry in mice carrying *Pacap-IRES-Cre* and/or *Trh-2a-Dre* following AAV injection. C) 8-hour food intake of PVH$^{Pacap/Trh}$-hM3Dq mice following injection of vehicle or CNO under *ad libitum* conditions in the light period. (N = 7 mice; mean of two trials per mouse; two-tailed paired t-test). **$p<0.01$.

**Figure S2. Generation and characterization of *Sim2-2a-Cre*, Related to Figure 1.**
A) Design of *Sim2-2a-Cre* allele. B) Sequence of *2a-Cre* insertion and homology arms. C) Example images of brain regions containing significant numbers of tdTomato-expressing neurons in a *Sim2-2a-Cre*::*tdTomato* mouse. D) Left: example images of tdTomato (magenta), *Sim2* (cyan), and *VGlut2* (blue) expression in the anterior PVH of a *Sim2-2a-Cre*::*tdTomato* mouse, as detected by smFISH against *Sim2* and *VGlut2* mRNA and endogenous tdTomato fluorescence. Arrowheads indicate cells coexpressing tdTomato, *Sim2*, and *VGlut2*. Right: quantifications of the % of tdTomato-expressing PVH neurons that express *Sim2* (left), % of *Sim2*-expressing PVH neurons that express tdTomato (middle), and % of tdTomato-expressing neurons that express *VGlut2* (right). (N = 3 mice, mean ± SEM).3V = third ventricle; 4V = fourth ventricle; III = oculomotor nucleus; cc = corpus callosum; cpd = cerebral peduncle; ILA = infralimbic area; int = internal capsule; DP = dorsal peduncular area; LV = lateral ventricle; MB = mammillary body; mtt = mammillothalamic tract; opt = optic tract; TT = taenia tecta; ZI = zona incerta.

**Figure S3. Generation and characterization of *Sim2-2a-FLPo*, Related to Figure 1.**
A) Design of *Sim2-2a-FLPo* allele. B) Sequence of *2a-FLPo* insertion and homology arms. C) Example images of brain regions containing significant numbers of *tdTomato*-expressing neurons in a *Sim2-2a-FLPo*::*tdTomato* mouse. D) Example image of tdTomato and *Sim2* expression in the anterior PVH of a *Sim2-2a-FLPo*::*tdTomato* mouse, as detected by smFISH against *Sim2* mRNA and endogenous tdTomato fluorescence. Arrowheads indicate cells coexpressing tdTomato and *Sim2*.3V = third ventricle; III = oculomotor nucleus; AC = anterior commissure; BNST = bed nucleus of the stria terminalis; cpd = cerebral peduncle; DMH = dorsomedial hypothalamus; LH = lateral hypothalamus; MB = mammillary body; mtt = mammillothalamic tract; opt = optic tract; VMH = ventromedial hypothalamus; ZI = zona incerta.

**Figure S4. Glutamatergic PVH$^{Sim2}$ neurons project to multiple brain regions, Related to Figure 1.**
A) Left: example images of tdTomato and fluorogold labelling in the anterior PVH of a *Sim2-2a-Cre*::*tdTomato* mouse following peripheral injection of fluorogold. Right:

quantification of the % of tdTomato-expressing PVH neurons that are labelled by fluorogold. (N = 4 mice, mean ± SEM). B) Representative images of mCherry expression in cell bodies (PVH, upper) and projection targets following injection of AAV-DIO-Syp-mCherry into the PVH of a *Sim2-2a-Cre* mouse. Inset: zoom of the ARC. C) Schematic of collateral mapping by injection of AAVretro-fDIO-Cre into one projection site (e.g. ARC) and AAV-DIO-Syp-mCherry into PVH, and subsequent examination of projections to another projection site (e.g. BNST). D) Representative images of Syp-mCherry expression in PVH, BNST, and ARC following AAVretro-fDIO-Cre injection into ARC (upper) or BNST (lower).3V = third ventricle; AC = anterior commissure; BNST = bed nucleus of the stria terminalis; LH = lateral hypothalamus; LS = lateral septum; opt = optic tract; MeA = medial amygdala; MHb = medial habenula; MPO = medial preoptic area; PVT = paraventricular thalamus.

**Figure S5. $ARC^{AgRP}$ neurons do not increase $PVH^{Sim2}$ activity; and ARC-projecting $PVH^{Sim2}$ neurons drive feeding, Related to Figure 1.**
A) Schematic of *in vivo* optogenetic stimulation of $ARC^{AgRP}$ terminals with simultaneous fiber photometry recording of $PVH^{Sim2}$ neuron activity. B) Mean ± SEM trace of $PVH^{Sim2}$ neuron activity before, during, and after 10s stimulation of $ARC^{AgRP}$ axons in PVH. (N = 4 mice; mean of 15 trials per mouse). C) Schematic of strategy for hM3Dq-mCherry expression in ARC-projecting $PVH^{Sim2}$ neurons. D) Representative images of resultant mCherry expression following injections into a *Sim2-2a-FLPo* (left) or wild-type (right) mouse. E) 3-hour food intake of mice expressing hM3Dq-mCherry in ARC-projecting $PVH^{Sim2}$ neurons following injection of vehicle or CNO. (N = 6 mice; two-tailed paired Wilcoxon signed-rank test). *$p<0.05$.

**Figure S6. $PVH^{Ucn3}$ neurons are not $ARC^{AgRP}$ neuron afferents, Related to Figure 1.**
A) Schematic of AAV-DIO-Syp-mCherry injection into the PVH of *Ucn3-Cre* mice. B) Representative images of resultant mCherry expression in the PVH (left) and mediobasal hypothalamus (right). C) Fraction of Npy-hrGFP-labelled ARC neurons that show EPSCs with monosynaptic latency following optogenetic activation of $PVH^{Ucn3}$ neuron projections by CRACM. D) Example recording from Npy-hrGFP neuron with optogenetic stimulation of $PVH^{Ucn3}$ neurons in the presence of artificial cerebrospinal fluid (aCSF). E) Schematic of AAV-DIO-hM3Dq-mCherry injection into the PVH of *Ucn3-Cre* mice. F) Representative image of resultant mCherry expression in PVH. G) Daytime (ZT3-6) food intake following injection of vehicle or CNO. (N = 8 mice, two-way ANOVA: no significant effect of treatment or interaction). ns = not significant.3V = third ventricle; ARC = arcuate nucleus; ME = median eminence; PVH = paraventricular hypothalamus; VMH = ventromedial hypothalamus.

**Figure S7. Overnight fasting increases Fos expression in $PVH^{Sim2}$ neurons, Related to Figure 3.**
A) Representative images of tdTomato and Fos expression in the anterior PVH of *Sim2-*

*2a-FLPo*::*tdTomato* mice following *ad libitum* food access or overnight fasting. Arrowheads indicate neurons coexpressing tdTomato and Fos. B) Percentage of tdTomato-labelled neurons expressing Fos (mean ± SEM). Each dot represents one mouse; total neurons across 4 sections spanning anterior-posterior PVH. N = 3 mice per condition; two-tailed unpaired t-test. **p<0.01.

**Figure S8. Ghrelin, insulin and leptin levels are unchanged following 1 hour of fasting, Related to Figure 3.**
A,B,C) Plasma levels of ghrelin (A), insulin (B), and leptin (C) at ZT12 or ZT13, either following *ad libitum* food access, or 1 hour of fasting (from ZT12 to ZT13). N = 3 - 6 mice per condition; one-way ANOVA. ns = not significant.

**Figure S9. Water deprivation does not activate PVH$^{Sim2}$ /or ARC$^{AgRP}$ neurons; and food removal does not affect DMH$^{Lepr}$ neuron activity, Related to Figure 3.**
A,C) Recording of ARC$^{AgRP}$ (A) or PVH$^{Sim2}$ (C) neuron activity in the water deprivation and *ad libitum* conditions (mean ± SEM over mice; 30-minute moving mean). Vertical lines indicate the beginning and end of water deprivation. (A: N = 6 mice; C: N = 10 mice). B,D) Mean activity in 1-hour bins between ZT11 and ZT16, for ARC$^{AgRP}$ (B) and PVH$^{Sim2}$ (D) neuron recordings. Lines indicate individual mice; box and whiskers indicate upper/lower quartiles and minimum/maximum data points, respectively. Outliers indicated by diamond. Two-way repeated measures ANOVA. E) Schematic of approach for fiber photometry recording from vDMH$^{Lepr}$ neurons. F) Representative image of GCaMP6s expression in vDMH$^{Lepr}$ neurons; dashed line indicates optic fiber placement. G) Recording of vDMH$^{Lepr}$ neuron activity in the fast and *ad libitum* conditions (mean ± SEM over mice; 30-minute moving mean). Vertical lines indicate the beginning and end of fasting. (N = 6 mice). H) Mean activity in 1-hour bins between ZT11 and ZT16 from vDMH$^{Lepr}$ recordings. Lines indicate individual mice; box and whiskers indicate upper/lower quartiles and minimum/maximum data points, respectively. Outliers indicated by diamond. Two-way repeated measures ANOVA. ns = not significant.

**Figure S10. PVH$^{Trh}$ neuron inhibition delays the response of ARC$^{AgRP}$ neurons to food removal, but does not affect ongoing ARC$^{AgRP}$ neuron activity in the fasted state, Related to Figure 3.**
A) Schematic of approach for fiber photometry recording from ARC$^{AgRP}$ neurons (using *Npy-IRES-FLPo*) with chemogenetic inhibition of PVH$^{Trh}$ neurons. B) Representative images of GCaMP6s and hM4Di-mCherry expression in PVH (left) and ARC (right). Dashed line indicates optic fiber placement. C) Recording of ARC$^{AgRP}$ neuron activity (mean ± SEM) in the fast condition with injection of vehicle or CNO (to inhibit PVH$^{Trh}$ neurons) at ZT11.5. Inset: magnification of activity between ZT11 and ZT15. (N = 10 mice). D) Time constant ($\tau$) of increase in ARC$^{AgRP}$ neuron activity for recordings shown in (C). Two-tailed paired t-test. E) Mean activity in 1-hour bins between ZT11 and ZT16 from recordings in (C). Two-way repeated measures ANOVA; post-hoc comparisons at

each time point with Holm-Sidak correction. F) Schematic of AAV-DIO-hM4Di-mCherry injection into the PVH of *Sim2-2a-Cre* mice. G) Daytime (ZT3-6) food intake following overnight fast, and injection of vehicle or CNO 1 hour prior to the start of refeeding. (N = 12 mice; two-way repeated measures ANOVA). ***p<0.001; *p<0.05; ns = not significant.For (D), (E), and (G), lines indicate individual mice; box and whiskers indicate upper/lower quartiles and minimum/maximum data points, respectively. Outliers indicated by symbol.

**Figure S11. PVH$^{Sim2}$ and ARC$^{AgRP}$ neuron activity during fasting peaks early in the dark period, Related to Figure 3.**
A) Schematic of experimental design: fasting is initiated at ZT12, ZT9 or ZT6. B,C) Recording of ARC$^{AgRP}$ (B) or PVH$^{Sim2}$ (C) neuron activity in the *ad libitum* condition or over fasting initiated at different times (mean ± SEM over mice; 30-minute moving mean). Colored vertical lines indicate the beginning and end of fasting for each condition. (B: N = 5 mice; C: N = 10 mice). D,E) Time of the highest level of ARC$^{AgRP}$ (D) or PVH$^{Sim2}$ (E) neuron activity shown in (B/C) for each mouse between ZT7 and ZT5. Boxplot: box and whiskers indicate upper/lower quartiles and minimum/maximum values, respectively. Symbols indicate outliers. One-way repeated measures ANOVA. ns = not significant.

**Figure S12. The choice of threshold does not affect the pattern of ARC$^{AgRP}$ and PVH$^{Sim2}$ neuron activity around pokes, Related to Figure 4.**
A,B) ARC$^{AgRP}$ (A) or PVH$^{Sim2}$ (B) neuron activity aligned to pokes (mean ± SEM over pokes). The calculation includes all pokes with no pokes in the preceding 45 (left), 30 (center-left), or 10 s (center-right), or all pokes (right). A: N = 7 mice B: N = 4 mice.

**Figure S13. During fasting, PVH$^{Sim2}$ neuron responses to unrewarded pokes gradually shift to earlier time points, Related to Figure 4.**
A,B) ARC$^{AgRP}$ (A) or PVH$^{Sim2}$ (B) neuron activity aligned to pokes, split by the decile of these pokes (as a percentage of the total pokes during fasting; mean ± SEM over pokes). A: N = 7 mice; B: N = 4 mice.

**Figure S14. PVH$^{Sim2}$ synaptic silencing reduces energy expenditure, Related to Figure 5.**
A) Body mass of male mice used for metabolic experiments (Figures 5F, 5G, and S14) at 4 weeks post-injection. (Mean ± SEM, N = 7 mice per condition; two-tailed unpaired t-test). B-D) Energy expenditure (B), cumulative distance travelled (C), and respiratory exchange ratio (D) over the 2-day indirect calorimetry experiment (mean ± SEM).

**Figure S15. PVH$^{Sim2}$ neurons express Cbln2 and Bdnf, Related to Figure 5.**
Example images of *Cbln2* (magenta) and *Sim2* (yellow, upper row) or *Bdnf* (magenta) and *Sim2* (yellow, lower row) expression in the anterior PVH, as detected by smFISH.

Arrowheads indicate PVH neurons coexpressing *Cbln2* and *Sim2* (upper row) or *Bdnf* and *Sim2* (lower row).
